# A complete-genome view of phylum Nanobdellota and recurrent Form III RuBisCO transfer between archaea and Patescibacteriota

**DOI:** 10.64898/2026.05.13.725050

**Authors:** Torben N. Nielsen, Lauren M. Lui

## Abstract

The archaeal phylum *Nanobdellota* (formerly *Nanoarchaeota*) was previously represented by four complete genomes. We present 208 complete *Nanobdellota* genomes from Oxford Nanopore metagenomes of the Baltic Sea water column and Fennoscandian groundwater (69–201 m below sea level), rotated to the ORC1/Cdc6 replication origin — a 52-fold expansion of complete-genome representation.

Across the ar53 supermatrix and a *Nanobdellota*-tuned 71-marker supermatrix on 1,239 taxa, the named GTDB orders within *Nanobdellota* are recovered as monophyletic clades, including the three orders that dominate our environmental sampling: *Woesearchaeales*, *Pacearchaeales*, and the GTDB placeholder order SCGC-AAA011-G17. This is consistent with the existing GTDB R232 order-level circumscription. We retire the SCGC-AAA011-G17 placeholder name, replacing it with a complete-genome-anchored SeqCode nomenclatural chain (*Maxwellarchaeales* ord. nov., *Maxwellarchaeaceae* fam. nov., *Maxwellarchaeum* gen. nov., and *Maxwellarchaeum balticum* sp. nov.) without altering the order-level circumscription.

*Pacearchaeales* and *Maxwellarchaeales* retain no central or energy metabolism beyond Form III RuBisCO, PEP synthase, and ferredoxin; *Woesearchaeales* retains partial glycolysis and a V/A-type ATPase. A 4,262-tip phylogeny of *rbcL* (the RuBisCO large-subunit gene) identifies nine candidate archaea-to-*Patescibacteriota* Form III RuBisCO transfer events — including one to a Baltic *Minisyncoccia* — versus two reciprocal candidates, consistent with archaea-to-CPR being the more frequently identified direction in our data. All 256 *Nanobdellota* genomes (208 complete + 48 high-quality non-circular), the ar71 marker set with its 1,239-taxon ML tree, 154 *Nanobdellota*-trained HMMs for KEGG-ortholog detection in DPANN proteomes (94 ROBUST), and the 4,262-tip rbcL reference tree are released as a community resource, alongside the full analysis archive — alignments, intermediate trees, structural predictions, and per-step scripts — at Zenodo (DOI 10.5281/zenodo.20174424; see *Using the resource*).

## Introduction

The phylum *Nanobdellota* — formerly known as *Nanoarchaeota* — was first encountered through the discovery of *Nanoarchaeum equitans*, a 491-kbp hyperthermophilic obligate symbiont of *Ignicoccus hospitalis* isolated from a submarine hydrothermal vent (Huber et al. 2002, Waters et al. 2003). *N. equitans* possessed the smallest archaeal genome then known and was the first archaeon demonstrated to depend entirely on a host for viability. The phylum was formally validated as *Nanobdellota* following the description of *Nanobdella aerobiophila*, a thermoacidophilic ectosymbiont (host-attached but not internalized) from a Japanese hot spring (Kato et al. 2022). GTDB recognizes *Nanobdellota* as a phylum within the DPANN superphylum, containing at least 12 named orders that span thermophilic, aquatic, and subsurface environments (Parks et al. 2022, Rinke et al. 2021). Three orders dominate the environmental aquatic and subsurface samples we surveyed here: *Woesearchaeales*, the metabolically broadest of the three and named after Carl Woese; *Pacearchaeales*, a metabolically reduced sister order named after Norman Pace; and the GTDB placeholder order SCGC-AAA011-G17, here proposed as *Maxwellarchaeales* ord. nov. (*Taxonomic proposal*). These three orders are the focus of the present analysis.

DPANN is the unranked clade originally comprising *D*iapherotrites, *P*arvarchaeota, *A*enigmarchaeota, *N*anoarchaeota, and *N*anohaloarchaeota, now extended to additional phyla. GTDB treats it as a superphylum of more than ten archaeal phyla, of which *Nanobdellota* is one. The *Patescibacteriota* (Candidate Phyla Radiation, CPR) are a bacterial superphylum of similarly small-genome, often host-associated lineages, and are sometimes referred to as the bacterial counterpart to DPANN; the parallel is functional rather than phylogenetic, and is relevant here because the same environments commonly co-host both groups. *Minisyncoccus archaeiphilus*, recently cultured by Nakajima et al. (2025), is the first cultured *Minisyncoccia* isolate and was shown by those authors to live as an obligate parasite of a methanogen — the first cultured CPR demonstrated to live as an archaeal parasite. Nakajima et al. (2025) propose *M. archaeiphilus* as the type species of *Minisyncoccaceae* / *Minisyncoccales* / *Minisyncoccia* / *Minisyncoccota*, the latter representing a phylum-rank elevation of the lineage formerly catalogued in *Candidatus* Patescibacteria / candidate phyla radiation. We retain the GTDB R226 *Patescibacteriota* phylum-level assignment for our co-sampled Baltic *Minisyncoccia* genome (B10_u4232805) for consistency with the rest of the analyses presented here, since the Nakajima reclassification has not yet been propagated to GTDB. Most known DPANN diversity is represented by metagenome-assembled genomes (MAGs): partial genome reconstructions binned from short-read or hybrid metagenome data and typically distributed across multiple contigs at varying completeness. Complete genomes are single, closed contigs spanning the entire chromosome; they provide a stronger basis for negative claims about gene content (e.g., “this genome does not encode the electron transport chain”) than do fragmented MAGs, where the apparent absence of a gene is ambiguous between true loss and assembly gap.

A clade is a group consisting of a common ancestor together with all of its descendants; i.e., a monophyletic group. A grouping that contains a common ancestor plus only some of its descendants is paraphyletic (a familiar example: birds are dinosaurs, so the order Reptilia is paraphyletic if treated as excluding birds); a grouping that combines descendants of multiple independent ancestors is polyphyletic. Formal Linnaean naming under the International Code of Nomenclature of Prokaryotes requires a cultivated type strain; the SeqCode (Hedlund et al. 2022) is a parallel nomenclature framework that permits formal naming of prokaryotic taxa from genome sequences alone and requires named taxa to be monophyletic, a necessary alternative for lineages such as *Nanobdellota* where most taxa have not been cultivated.

DPANN archaea are characterized by small genomes, limited metabolic capacity, and frequent symbiotic or parasitic lifestyles (Castelle et al. 2018, Dombrowski et al. 2019, He et al. 2021, Huang and Spang 2025). Diverse interaction modes have been described across DPANN, including ectosymbiosis via nanotubes (Johnson et al. 2024) and host-dependent replication in *Nanohaloarchaeota* (Hamm et al. 2019). Form III RuBisCO — a large-subunit-only enzyme that functions in nucleotide salvage rather than carbon fixation — has been reported in DPANN archaea via lateral gene transfer (Jaffe et al. 2019), but its distribution across *Nanobdellota* orders and its relationship to other metabolic capabilities has not been characterized at scale with complete genomes. Baker et al. (2025) recently proposed that DPANN originated from free-living euryarchaeal ancestors, implying that the small genomes and host dependency of extant DPANN are derived rather than ancestral traits (i.e., evolved later within DPANN from a larger-genome free-living ancestor, rather than inherited as DPANN’s starting condition). Whether the metabolic diversity within *Nanobdellota* reflects different stages along a reductive path or independent divergent adaptations to the same ecological constraints is unknown.

Prior to this work, only four complete genomes existed for the entire phylum *Nanobdellota*: *Nanoarchaeum equitans* Kin4-M (491 kbp), *Nanobdella aerobiophila* MJ1 (669 kbp), *Candidatus* Nanopusillus acidilobi (606 kbp), and *Nanobdellota* archaeon YN1 (773 kbp) — all from thermophilic or hyperthermophilic environments. GTDB R232 also contains one additional single-contig high-quality MAG (GCA_028279745.1, ∼1.3 Mbp, 100% CheckM2-complete) that is not closed at the assembler level. The remaining public *Nanobdellota* assemblies are fragmented metagenome-assembled genomes (MAGs; bin-level reconstructions assembled from short or hybrid metagenome data) whose incompleteness prevents reliable distinction between genuine metabolic absences and assembly artifacts. At the genome sizes (553 kbp to 2,676 kbp) and ∼90% coding density of *Nanobdellota*, every gene counts: a missing gene in a fragmented MAG is ambiguous, but a missing gene in a complete genome is a genuine absence.

Here we present 208 complete *Nanobdellota* genomes from Oxford Nanopore metagenomes of two regions: the Baltic Sea water column (four samples) and Fennoscandian groundwater (two sites at 69 and 201 m below sea level, at the Äspö Hard Rock Laboratory underground research facility managed by the Swedish Nuclear Fuel and Waste Management Company, SKB). We combine the 208 with 238 high-quality NCBI MAGs spanning 12 named orders to construct a 446-genome phylum-level comparative analysis. A further 48 high-quality single-contig but non-circular *Nanobdellota* assemblies from the same metagenomes are released alongside as a community resource but are not used in the phylogenetic analyses, which restrict to assembler-confirmed circular topology. Two questions organise the analysis. First, is the GTDB order-level taxonomy of *Nanobdellota* consistent with phylogeny when complete-genome substrate is brought to bear under multiple inference methods, and if not, what taxonomic revision is warranted under the SeqCode? Second, does the metabolic gradient across *Nanobdellota* orders, together with the distribution of Form III RuBisCO across the phylum, show evidence of lateral exchange with co-sampled *Patescibacteriota*, and if so, in what direction? Subsidiary descriptive results across the resource (species-level diversity and biogeography across the three environments; placement of environmental versus thermophilic orders; phylum-wide single-copy core and a *Nanobdellota*-specific phylogenomic marker set; per-order metabolic profiles and genome-size predictors of metabolic complexity) are reported alongside. On the first question, the GTDB order-level taxonomy of *Nanobdellota* is consistent with our complete-genome ar53 and ar71 phylogenies: every named multi-tip order, including the three environmental orders, is recovered as a monophyletic clade, and no taxonomic revision is warranted at order rank. We provide a species-rank SeqCode protologue for a representative complete genome of one of those orders. On the second question, the rbcL gene tree identifies recurrent archaea-to-CPR Form III RuBisCO transfer as a feature of the same small-genome lineages that drive the within-phylum metabolic divide.

## Results

### Recovery of 256 Nanobdellota genomes from the Baltic Sea water column and Fennoscandian groundwater

A total of 256 high-quality *Nanobdellota* genomes were recovered from Oxford Nanopore metagenomes: 208 complete from two regions (the Baltic Sea water column, four samples; Fennoscandian groundwater, two samples at 69 and 201 m below sea level) and 48 additional high-quality non-circular genomes. Of the 208 complete genomes, 123 are from the Baltic Sea, 29 from KR0015B (Äspö Hard Rock Laboratory, Sweden; 69 m below sea level), and 56 from SA1420A (Äspö Hard Rock Laboratory, Sweden; 201 m below sea level). GTDB-Tk classification assigned all 256 genomes to the same three orders across all environments: *Woesearchaeales* (123 complete: 77 Baltic, 13 KR0015B, 33 SA1420A), *Pacearchaeales* (54 complete: 29 Baltic, 10 KR0015B, 15 SA1420A), and the GTDB R232 placeholder order *SCGC-AAA011-G17* (31 complete: 17 Baltic, 6 KR0015B, 8 SA1420A; *SCGC* carries its origin as a Single-Cell-Genomics-Center placeholder; Rinke et al. 2021). The placeholder *SCGC-AAA011-G17* clade is here proposed as *Maxwellarchaeales* ord. nov. (*Taxonomic proposal*); we use the proposed name *Maxwellarchaeales* throughout the Results, Tables, and Discussion of this manuscript and retain *SCGC-AAA011-G17* in Methods only where we are factually documenting GTDB-Tk output. Genome sizes range from 553 kbp to 2,676 kbp, with *Woesearchaeales* having the largest genomes (median 1,409 kbp) and *Pacearchaeales* the smallest (median 739 kbp). Gene prediction on the 208 complete genomes yielded 274,837 protein-coding genes (mean 1,321 per genome). The complete genomes were processed for rotation to ORC1/ Cdc6, the archaeal replication initiation protein (an orthologue of eukaryotic ORC1 and Cdc6). ORC1/Cdc6 (K10725) is present at strict KofamScan threshold in 206 of 208 complete genomes; the two exceptions (`KR0015B_u14002131`, `SA1420A_u36387404`; flagged DEMOTED_no_ORC1 in `genome_flags.tsv`) carry only sub-threshold K10725 hits, consistent with frameshifted or otherwise non-canonical ORC1 (Methods *Rotation to ORC1/Cdc6*). The bacterial dnaA (K02313) is absent throughout (zero strict hits; sub-threshold K02313 hits were re-screened against UniRef90 by MMseqs2 easy-search and found to be CDC48 AAA+ ATPases — a different AAA+ family — rather than archaeal dnaA orthologs).

### 238 high-quality NCBI Nanobdellota genomes span at least 12 orders

To place the Baltic genomes in a global context, we downloaded all 870 *Nanobdellota* assemblies from NCBI (after GCA/GCF deduplication). CheckM2 identified 245 as high-quality (≥ 90% completeness, < 5% contamination). GTDB-Tk classification confirmed 238 of these as *Nanobdellota*, with the remaining 7 assigned to other phyla. The 238 *Nanobdellota* genomes span at least 12 orders (3 additional genomes are unclassified to order level): *Woesearchaeales* (82), *Maxwellarchaeales* (44), *Pacearchaeales* (29), *Nanobdellales* (16), *Jingweiarchaeales* (12), *Parvarchaeales* (11), JAPDLS01 (11), CSSed11-243R1 (9), UBA10117 (7), *Tiddalikarchaeales* (7), WJKC01 (6), and JAQUSD01 (1). The combined dataset of 446 genomes (208 complete from this study + 238 high-quality NCBI MAGs) is the largest complete-genome-anchored survey of the phylum to date (Figure 2A).

### 145 species from 208 complete genomes: no shared species between environments

Species-level clustering of the 208 complete genomes at 95% ANI identified 145 species: 68 Baltic-only, 25 KR0015B-only, 49 SA1420A-only, and 3 species shared between KR0015B and SA1420A (Figure 2B). Not a single species is shared between the Baltic Sea and either KR0015B or SA1420A, despite both environments harboring the same three orders. The three shared KR0015B–SA1420A species are consistent with the physical proximity of the two Äspö sites. Zero species overlap with the 238 NCBI genomes. The closest Baltic–NCBI match is 85.1% ANI with < 20% alignment fraction, well below the species boundary. The only NCBI genomes with any detectable ANI to our genomes are from Olkiluoto nuclear waste repository in Finland (367–384 m depth; GCA_018901245.1) and the Äspö Hard Rock Laboratory in Sweden (201–448 m depth; GCA_018819765.1, GCA_043823355.1). Within each environment, zero cross-order ANI is detectable — the three orders are fully isolated at the nucleotide level.

### Phylogenomic structure: thermophilic orders basal, environmental orders derived

Two phylogenies were inferred. The ar53 phylogeny was inferred from a 446-genome alignment (208 complete from this study + 238 NCBI) of 53 archaeal marker genes (ar53, a set of single-copy genes conserved across Archaea and used by GTDB for taxonomic placement; 8,062 alignment positions after masking), yielding a 444-tip ML tree after IQ-TREE collapsed two identical sequences during inference. The inference used IQ-TREE v3.0.1 under the LG+F+R10 model, midpoint-rooted, with 1,000 ultrafast bootstrap (UFBoot — IQ-TREE’s bootstrap-resampling-based branch support that scales to large alignments; Hoang et al. 2018) replicates and 1,000 SH-aLRT (Shimodaira–Hasegawa-like approximate likelihood-ratio test; Guindon et al. 2010) replicates. The ar71 phylogeny was inferred from a 71-marker *Nanobdellota*-tuned supermatrix on 1,239 taxa drawn from our complete-genome set and 1,031 GTDB R232 high-quality *Nanobdellota* MAGs, under a single LG+F+R10 model (Methods *ar71 phylogeny*). All named orders with two or more representatives are recovered as monophyletic clades in both trees, including *Woesearchaeales* itself (205 tips at the 444-tip ar53 scale, 728 at the 1,239-tip ar71 scale; Discussion *Woesearchaeales is recovered monophyletic*). The ar71 tree additionally resolves *Parvarchaeales* (58 tips) and JAPDLS01 (10 tips) as monophyletic, both of which were non-monophyletic at the 444-tip ar53 scale. Bootstrap support is strong on both trees: on the ar53 tree, 79.5% of internal nodes have UFBoot ≥ 95 and 91.3% have UFBoot ≥ 80 (Figure 1); on the ar71 tree, 85.4% have UFBoot ≥ 95 and 91.7% have UFBoot ≥ 80.

**Figure 1.**
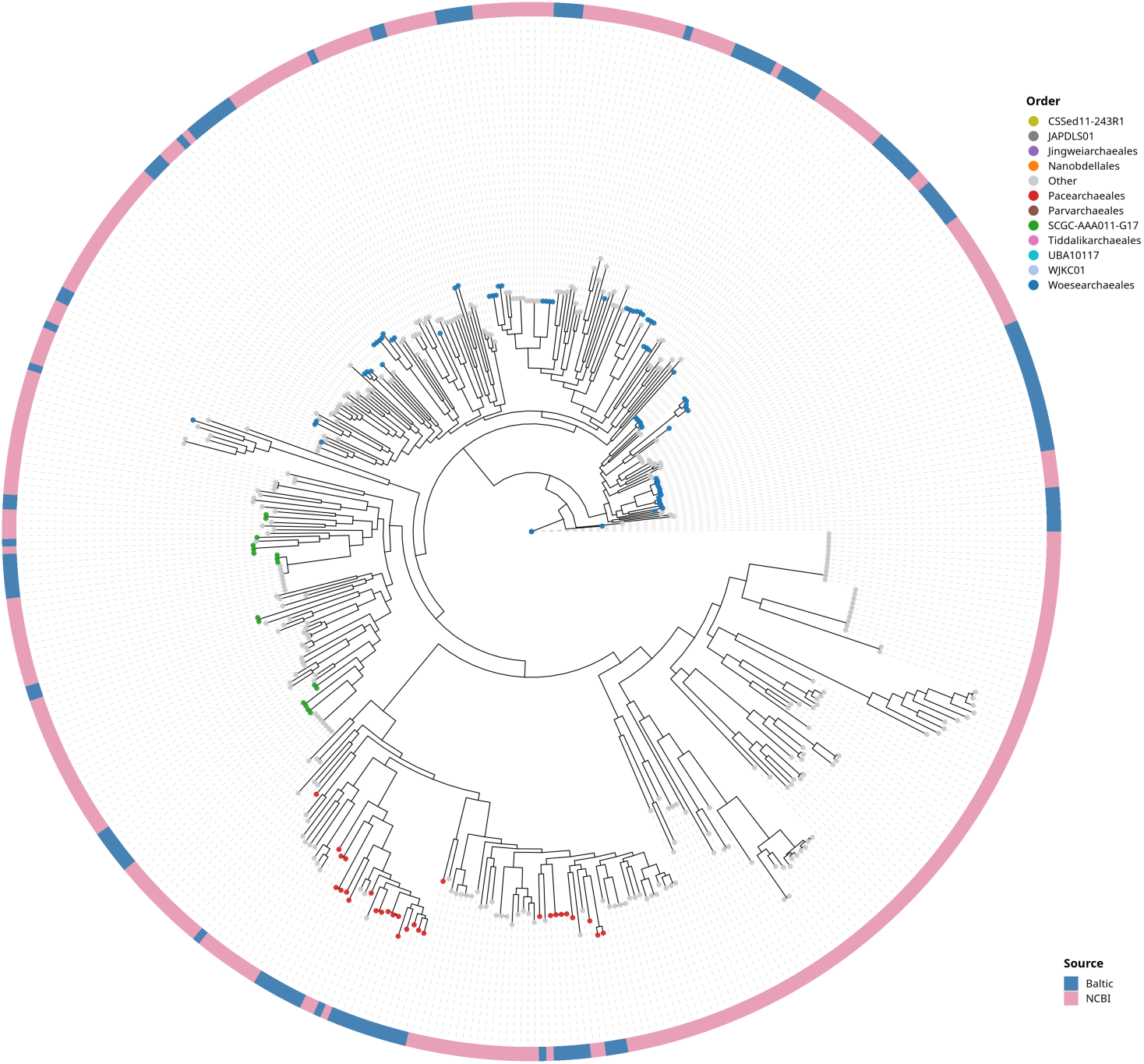
Radial ar53 supermatrix phylogeny of *Nanobdellota* (444-tip ML tree from a 446-genome alignment: 208 complete from this study + 238 NCBI; two identical sequences were collapsed by IQ-TREE before inference). Inferred from 53 archaeal marker genes (ar53; 8,062 positions) under the LG+F+R10 model with 1,000 ultrafast bootstrap replicates. Branches are colored by GTDB order (see legend; the order list also includes the basal-clade orders UBA10117, JAPDLS01, and JAQUSD01 that are present in the alignment but appear as small contributing groups of tips at this rendering scale). The outer ring indicates genome source: steel blue = Baltic Sea, dark blue = Fennoscandian groundwater, rose = NCBI. The tree is midpoint-rooted; thermophilic orders (*Nanobdellales*, *Jingweiarchaeales*, *Parvarchaeales*, *Tiddalikarchaeales*, CSSed11-243R1, WJKC01, JAPDLS01, JAQUSD01) are basal and environmental aquatic orders (*Woesearchaeales*, *Pacearchaeales*, *SCGC-AAA011-G17* — here renamed *Maxwellarchaeales*; *Taxonomic proposal* — and UBA10117) are derived. Each named multi-tip *Nanobdellota* order, including *Woesearchaeales*, forms a single bipartition in the inferred tree (Methods *Per-order monophyly assessment*; Discussion *Woesearchaeales is recovered monophyletic*). 79.5% of internal nodes have UFBoot support ≥ 95.

**Figure 2.**
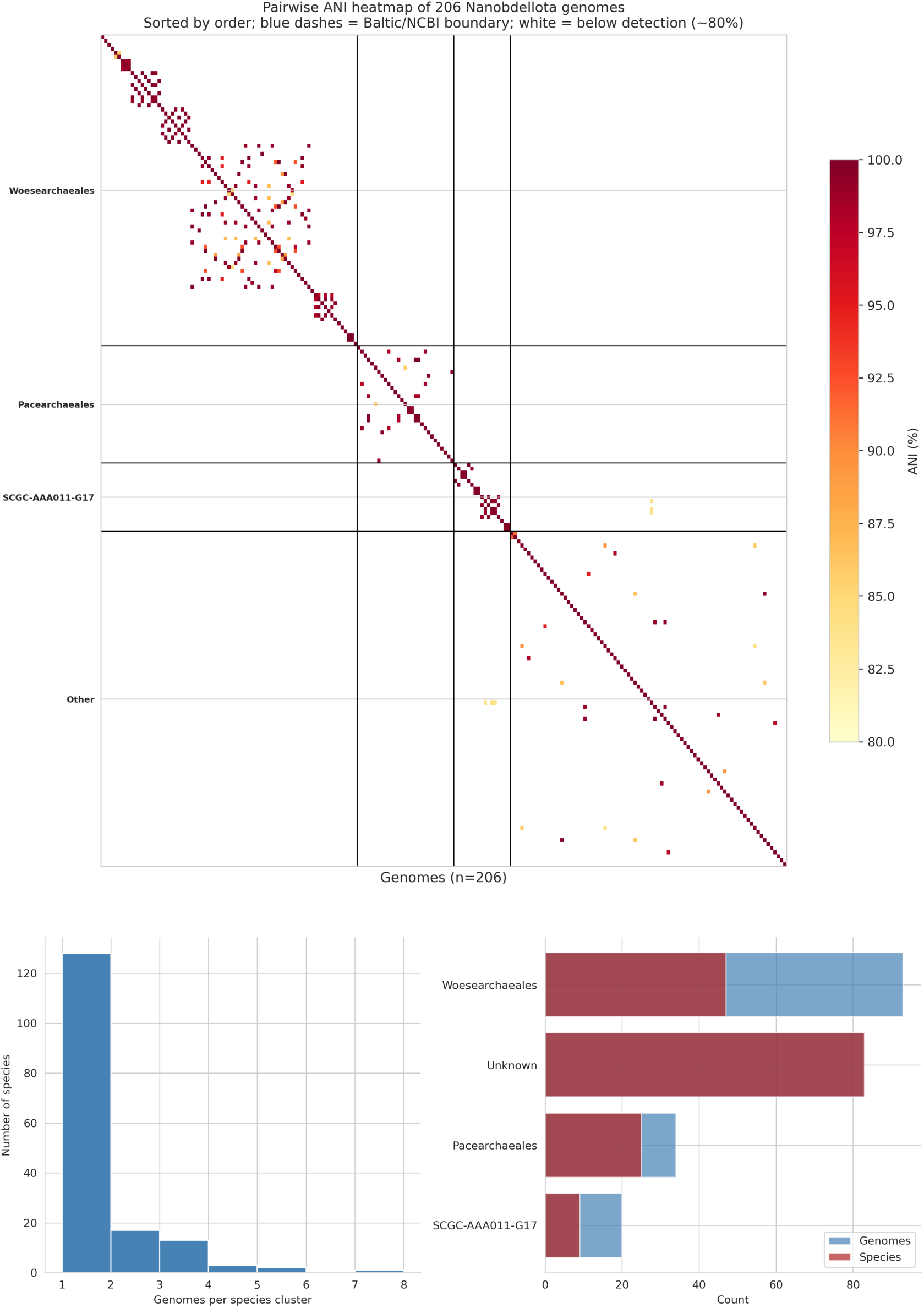
(A) Pairwise ANI heatmap of 208 complete *Nanobdellota* genomes sorted by order. Colored bars indicate environment: steel blue = Baltic Sea, dark blue = KR0015B, navy = SA1420A. White space indicates ANI below the detection limit (∼80%). The block-diagonal structure reflects within-order species clusters with zero cross-order ANI. (B) Bar chart of species diversity: 145 species at 95% ANI from 208 complete genomes, colored by environment. 68 Baltic-only, 25 KR0015B-only, 49 SA1420A-only, 3 shared between Äspö sites, zero shared with Baltic or NCBI.

Midpoint rooting places the thermophilic and extreme-environment orders in a basal clade: *Nanobdellales* (hot springs), *Jingweiarchaeales* (hot spring sediments), *Parvarchaeales* (acid mine drainage), *Tiddalikarchaeales* (hydrothermal deposits and landfill leachate), CSSed11-243R1, WJKC01 (deep-sea hydrothermal deposits), JAPDLS01 (hot spring sediments), JAQUSD01, and 3 unclassified genomes. The three environmental aquatic orders (*Woesearchaeales*, *Pacearchaeales*, and *Maxwellarchaeales*) together with UBA10117 (7 genomes) form a derived clade. Midpoint rooting assumes approximately clock-like evolution; the basal placement of thermophilic orders should be interpreted cautiously given the potentially elevated substitution rates in these lineages.

### RuBisCO and the metabolic divide across Nanobdellota orders

RuBisCO (ribulose-1,5-bisphosphate carboxylase/oxygenase) exists in multiple forms (Forms I, II, III, and IV are the canonical Linnaean form-numbering of RuBisCO families based on subunit composition and substrate specificity). Forms I and II function in the Calvin cycle for carbon fixation; Form III, characteristic of archaea but also present in select bacterial phyla, lacks the small subunit (rbcS) and instead functions in a nucleotide salvage pathway that recycles AMP via ribulose-1,5-bisphosphate (Sato et al. 2007). Throughout this paper we detect candidate rbcL by KEGG ortholog K01601 (the KEGG large-subunit ortholog, which spans Forms I, II, III, and IV at the gene level) and confirm Form III identity by absence of the small subunit rbcS (K01602; verified at both sequence and structural level via KofamScan and Foldseek search against AlphaFold-DB rbcS references; Methods *rbcS structural absence check*) together with absence of a sequence-level ortholog call for phosphoribulokinase PRK (K00855) — the two enzymes that distinguish Form I/II Calvin-cycle RuBisCO from Form III nucleotide-salvage RuBisCO. The KEGG ortholog database assigns each functional protein family an opaque “K-number” identifier, against which proteomes are searched with KofamScan KEGG HMMs; KO numbers throughout this paper are KEGG ortholog identifiers.

The large subunit of RuBisCO (rbcL, K01601) is present at strict KofamScan threshold in our complete-genome set at 100% of *Pacearchaeales* (54/54), 87% of *Maxwellarchaeales* (27/31), and 11% of *Woesearchaeales* (14/123) (Figure 3). The same pattern holds in both environments: *Pacearchaeales* has rbcL in all Baltic, KR0015B, and SA1420A complete genomes; *Woesearchaeales* has rbcL in < 12% regardless of environment. No genome in any order encodes the small subunit (rbcS, K01602; absent at both KofamScan strict threshold and Foldseek structural search against four AlphaFold-DB rbcS references), and no genome carries a KofamScan ortholog call for phosphoribulokinase (PRK, K00855), confirming Form III identity and ruling out the Calvin cycle. KofamScan on the 238 NCBI genomes recovers a lower per-order prevalence consistent with MAG incompleteness rather than genuine absence: 69% of *Pacearchaeales* (20/29), 77% of *Maxwellarchaeales* (34/44), and 10% of *Woesearchaeales* (8/82) (Figure 3, right panel). Pooled across the 446-genome combined set the per-order prevalences are 89% in *Pacearchaeales* (74/83), 81% in *Maxwellarchaeales* (61/75), and 11% in *Woesearchaeales* (22/205); the order-specific pattern is preserved at every level of the comparison (complete-only, NCBI-only, combined). Among the NCBI-only orders not represented in our complete-genome set, *Jingweiarchaeales* (100%, 12/12), CSSed11-243R1 (100%, 9/9), *Tiddalikarchaeales* (100%, 7/7), WJKC01 (100%, 6/6), and UBA10117 (71%, 5/7) have high prevalence, while *Nanobdellales* (6%, 1/16) and *Parvarchaeales* (9%, 1/11) have low prevalence.

**Figure 3.**
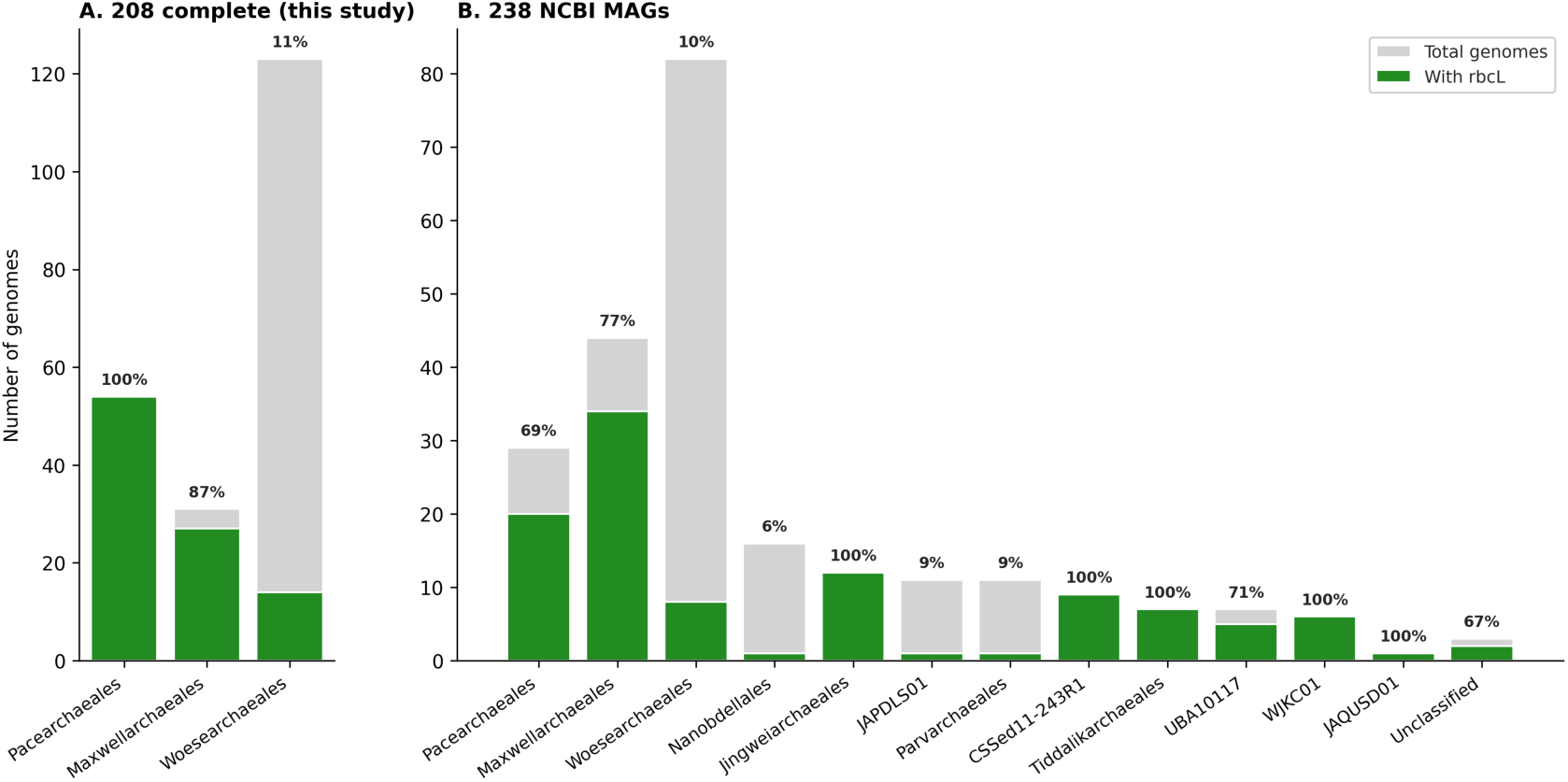
RuBisCO large subunit (rbcL) prevalence by order. Two panels: 208 complete genomes from this study (left) and 238 NCBI MAGs (right). Grey bars: total genomes per order; green bars: genomes with rbcL. Pooled 446-genome combined rates are tabulated below the figure. Per-order rates:

The three orders exhibit fundamentally different metabolic capabilities beyond RuBisCO, validated by KofamScan annotations cross-checked against UniRef90 (a clustered protein database at 90% identity from UniProt) best-hit descriptions for all 208 complete genomes. The non-RuBisCO metabolic content of the three orders centres on three molecules: phosphoenolpyruvate (PEP, a high-energy glycolytic intermediate); pyruvate, the canonical glycolytic endpoint; and ferredoxin, a small iron-sulfur cluster electron-carrier protein used in archaeal anaerobic energy metabolism. Two enzyme complexes connect these: PEP synthase (interconverts PEP and pyruvate at the cost or gain of one ATP, depending on direction; in the canonical forward direction PEP synthase consumes ∼2 ATP equivalents to make PEP from pyruvate, and the reverse reaction recovers approximately one ATP per cycle via the direct reverse reaction PEP + AMP + Pi → pyruvate + ATP), and pyruvate:ferredoxin oxidoreductase (PFOR, a four-subunit complex that oxidises pyruvate to acetyl-CoA and CO₂ while reducing ferredoxin; the [4Fe-4S] clusters — four iron and four sulfur atoms each in cubane-like geometry — that mediate electron transfer reside on the α, β, and δ subunits in the canonical *Pyrococcus*-paradigm assembly, with δ acting as a small ferredoxin-like electron-transfer subunit, while the γ subunit carries no metal centres and contributes a structural / TPP-pocket-stabilizing role rather than a redox role).

*Woesearchaeales* (123 complete) is the metabolically broadest order. The bulk of the glycolytic pathway is present in most genomes: glucokinase (K00845, 24/123), fructose-bisphosphate aldolase class II (K01624, 67/123), GAPDH (K00134, 117/123), phosphoglycerate kinase (K00927, 123/123), enolase (K01689, 122/123), and pyruvate kinase (K00873, 39/123). These six glycolytic KOs define the glycolytic gene count used below. V/A-type ATPase subunits, a proton pump distinct from the F-type ATP synthase used in oxidative phosphorylation, are present in approximately 40% of genomes (K02117/K02118/K02119, 47–50/123) — the only ATPase in any order. The conserved Pace/SCGC PEP-synthase / ferredoxin module is also retained at high frequency in *Woesearchaeales*: PEP synthase (K01007, 111/123) and ferredoxin (K05337, 113/123) are both present in ≈90% of *Woesearchaeales* genomes, so the module is order-non-specific within *Nanobdellota*.

PFOR α/β/δ/γ subunits (K00169–K00172) are detected in 33/31/30/29 of 123 *Woesearchaeales* (≈25%), a higher frequency than the 3/54 partial-PFOR *Pacearchaeales* but lower than the 17/31 *Maxwellarchaeales*.

Amino acid biosynthesis, riboflavin biosynthesis (6,7-dimethyl-8-ribityllumazine synthase K00794, 31/123; riboflavin pathway, KEGG map00740, not thiamine or biotin), and glycogen synthase (K00693, 68/123, 55%) are found predominantly or exclusively in *Woesearchaeales*.

*Pacearchaeales* (54 complete) is metabolically reduced. Beyond the universal information-processing core, *Pacearchaeales* genomes consistently encode RuBisCO (54/54), PEP synthase (K01007, 54/54), and ferredoxin (K05337, 47/54). PEP synthase is present in *Pacearchaeales* at 100%, and at ∼90% in *Woesearchaeales* (111/123) and *Maxwellarchaeales* (28/31), so the enzyme is order-non-specific within *Nanobdellota*. *Pacearchaeales* lacks all detectable upstream glycolytic enzymes that would provide pyruvate as the canonical forward-direction substrate, leaving the in vivo direction of PEP synthase in this order’s metabolic context uncharacterized by gene content alone. The full gene complement encodes no autonomous ATP-generating apparatus, but whether *Pacearchaeales* lineages live as obligate ectosymbionts, syntrophs, or under some other mode cannot be discriminated from gene content alone. A minority of genomes retain scattered glycolytic enzymes (GAPDH K00134 13/54, phosphoglycerate kinase K00927 13/54, pyruvate kinase K00873 9/54), but no genome encodes a majority of the pathway. Pyruvate:ferredoxin oxidoreductase subunits are detected in only 3/54 *Pacearchaeales* (all KR0015B; α, β, and δ subunits at strict KofamScan threshold; γ subunit absent across all 54). The three redox-bearing subunits (α, β, δ — the last being the ferredoxin-like [4Fe-4S]-bearing electron-transfer subunit) are present in those 3 genomes; no *Pacearchaeales* genome encodes the complete canonical four-subunit PFOR assembly. The catalytic significance of γ absence is uncertain since γ has no metal centres and contributes a structural / TPP-pocket-stabilizing role rather than a redox role. Succinyl-CoA synthetase α-subunit (K01902) is absent throughout (0/54).

*Maxwellarchaeales* (31 complete) resembles *Pacearchaeales*: RuBisCO (27/31), PEP synthase (K01007, 28/31), and ferredoxin (K05337, 30/31). Distinguishing features versus *Pacearchaeales* are pyruvate:ferredoxin oxidoreductase subunits (K00169 alpha 17/31, K00170 beta 13/31, K00171 delta 13/31, K00172 gamma 11/31) and succinyl-CoA synthetase α-subunit (K01902, 6/31), both retained in a substantial subset of *Maxwellarchaeales* but rare or absent across the 54 *Pacearchaeales* (Table 1).

**Table 1.**
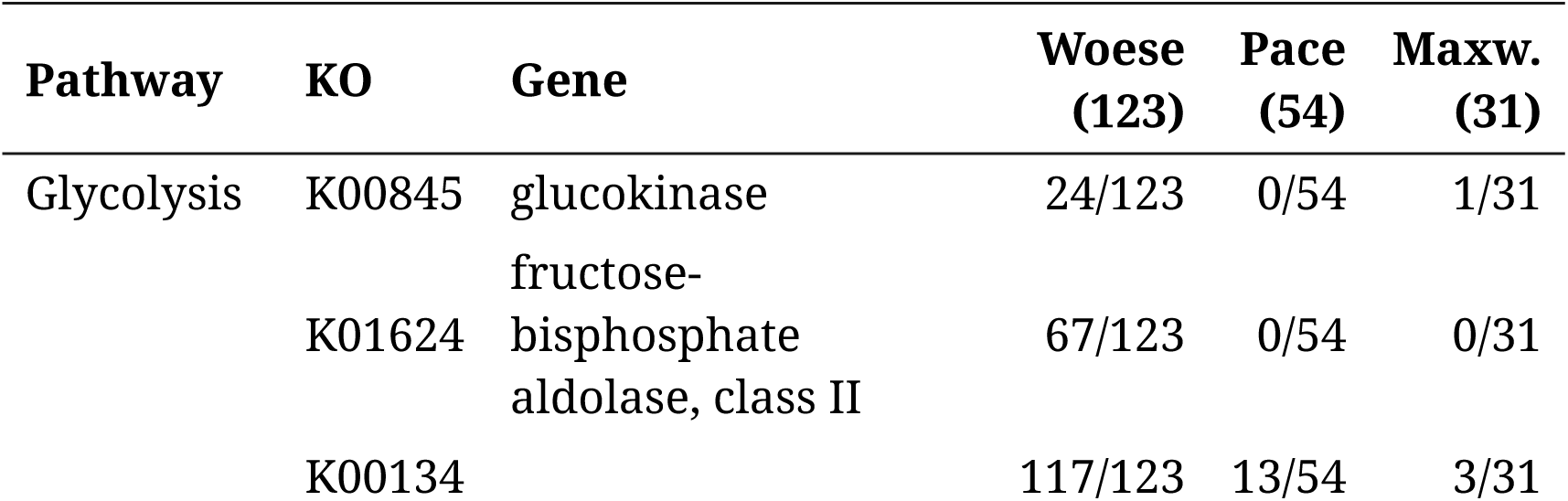
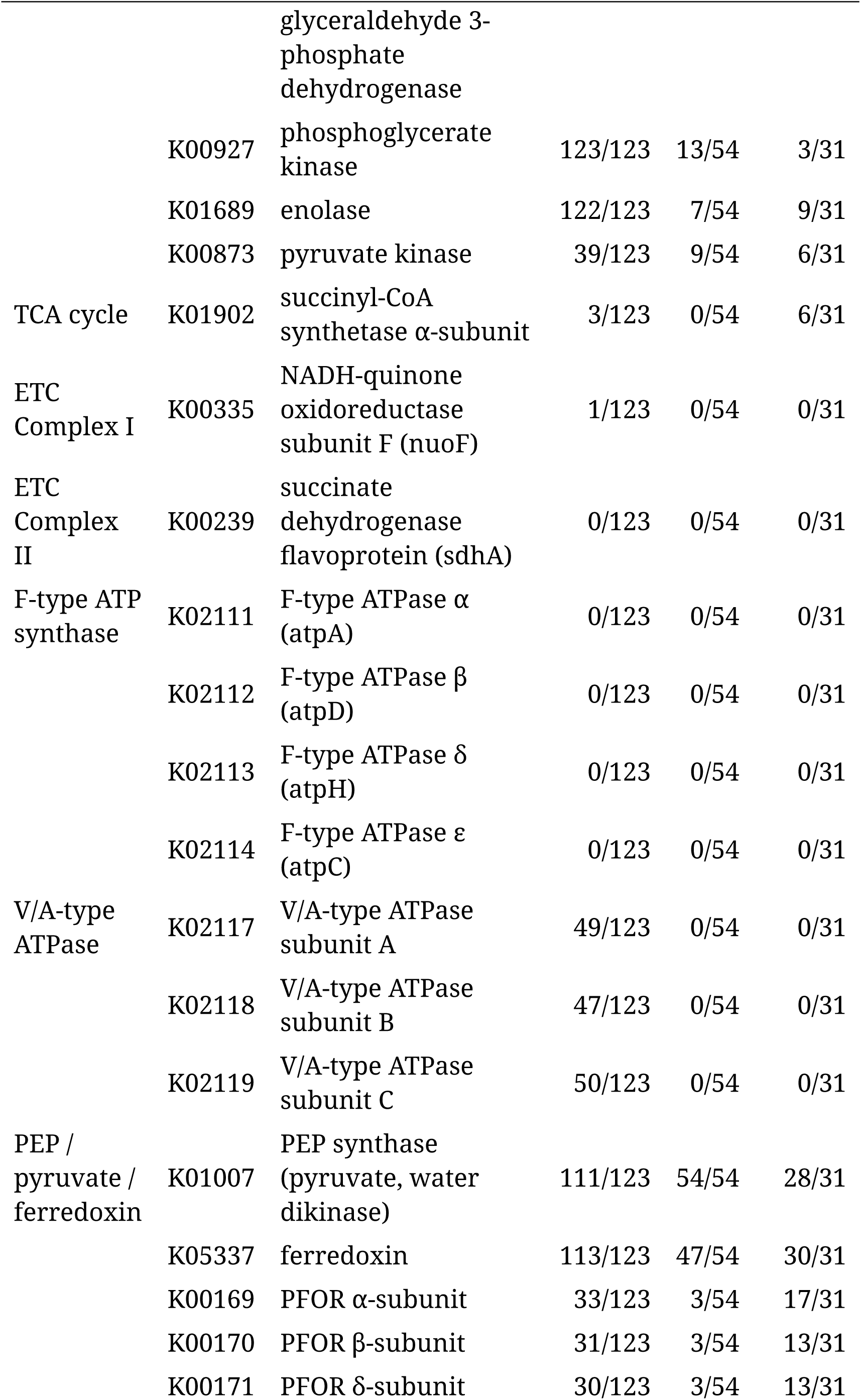
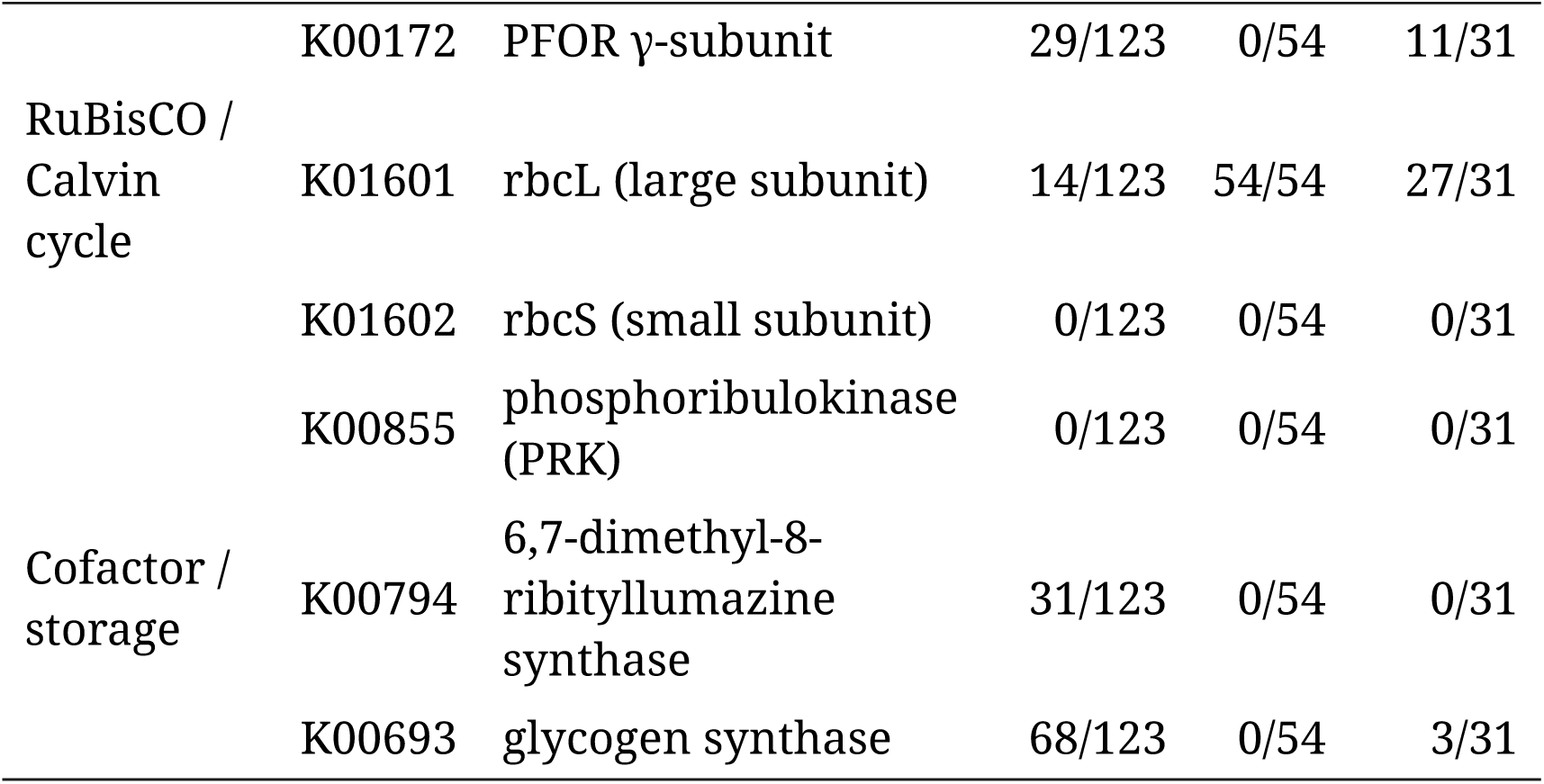
Per-order metabolic summary for 208 complete genomes. Columns: *Woesearchaeales* (123 genomes), *Pacearchaeales* (54 genomes), *Maxwellarchaeales* (31 genomes; here proposed as the SeqCode replacement for the GTDB R232 placeholder order *SCGC-AAA011-G17*; column abbreviated *Maxw.* below). Rows: key KOs across glycolysis, TCA cycle, electron transport chain, ATPase, RuBisCO, PEP/ pyruvate/ferredoxin metabolism, and cofactor/storage. Values: number of genomes with strict KofamScan hit / total. KO-to-gene mapping loaded canonically from /Kittens/Data/kofamscan/ko_list. Reproduced by scripts/build_table1.py, which writes scripts/ table1_per_order_metabolic.tsv.

At the per-genome level, RuBisCO prevalence anti-correlates with glycolytic gene count (Pearson r =-0.577, p < 10⁻³², n = 446 genomes with KofamScan data: 208 complete + 238 NCBI). However, this correlation is partly structured by order-level divergence — the orders themselves differ in both traits — and should not be interpreted as evidence of a within-lineage functional trade-off without further analysis. Genome size predicts the number of unique KOs per genome (Pearson r = 0.869, p < 10⁻⁹⁹, n = 446) (Figure 4).

**Figure 4.**
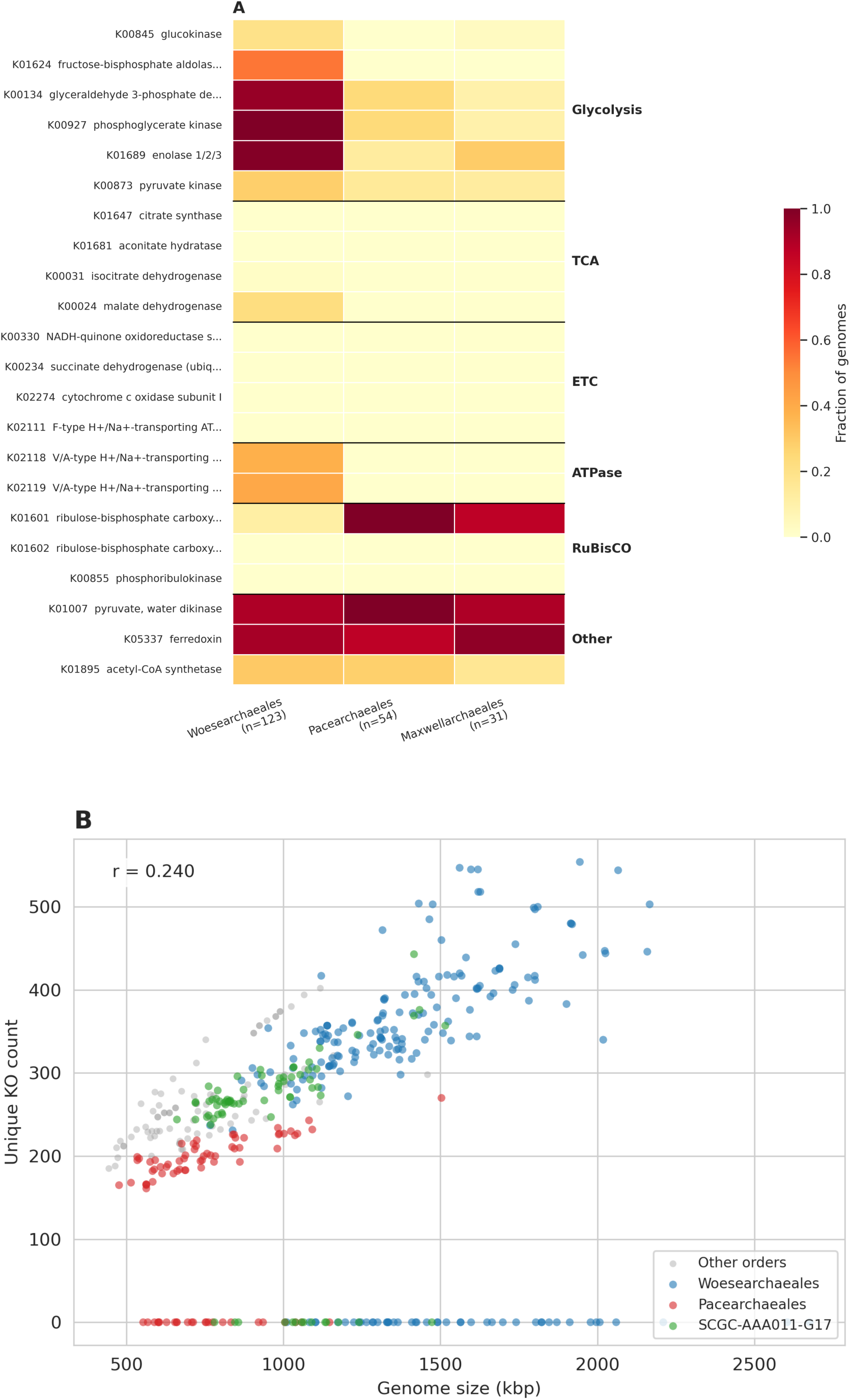
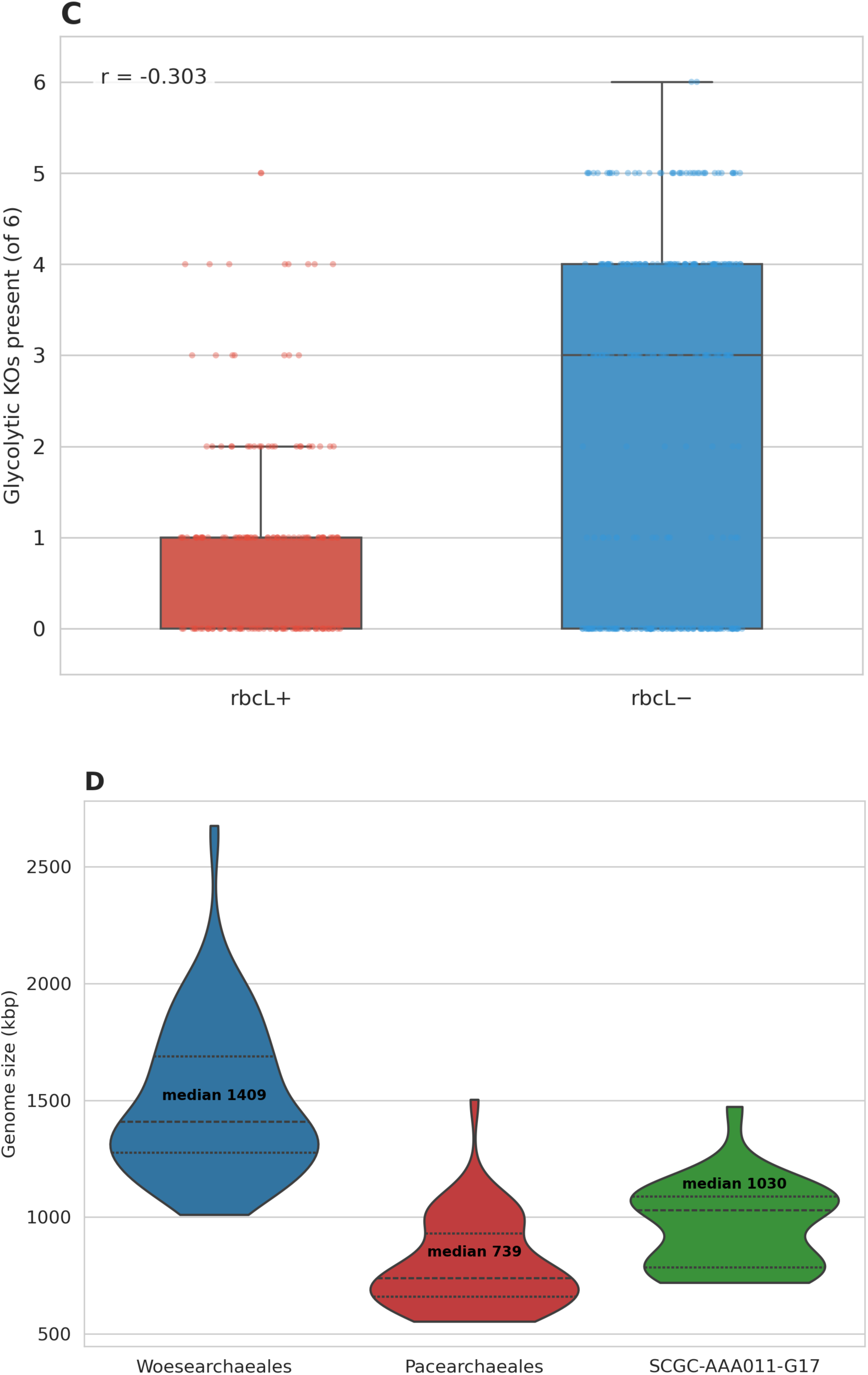
Per-order metabolic and genome-size structure across our 208 complete genomes (panels A and D) and the 446-genome combined cohort of 208 complete + 238 NCBI MAGs (panels B and C). (A) Heatmap of metabolic-pathway prevalence across the three principal orders represented in the complete-genome set (*Woesearchaeales* 123, *Pacearchaeales* 54, *Maxwellarchaeales* 31; *Maxwellarchaeales* shown in this paper as the proposed name for the GTDB R232 placeholder order *SCGC-AAA011-G17*). Rows: key metabolic functions selected across glycolysis, TCA, ETC, V/A-type ATPase, RuBisCO, ferredoxin, amino-acid and cofactor biosynthesis; columns: orders. Color intensity encodes the fraction of genomes per order with at least one strict KofamScan KEGG-HMM hit (deeper color = higher prevalence). (B) Per-genome scatter of genome size (kbp) versus number of unique KOs (KEGG ortholog identifiers) per genome (Pearson r = 0.869, p < 10⁻⁹⁹, n = 446 = 208 complete + 238 NCBI MAGs). Points coloured by order. The strong size–KO correlation reflects the per-genome metabolic-complexity gradient that the box plot in (C) decomposes by RuBisCO presence. (C) Box plot of the 6-gene glycolytic count (glucokinase + fructose-bisphosphate aldolase + GAPDH + PGK + enolase + pyruvate kinase) in rbcL-positive vs rbcL-negative genomes, illustrating the between-order RuBisCO–glycolysis anti-correlation (Pearson r = −0.577, n = 446). (D) Violin plot of genome-size distribution by order across the 208 complete genomes only (NCBI MAG sizes excluded so size distributions are not perturbed by MAG incompleteness): *Woesearchaeales* median 1,409 kbp ≈ 2× *Pacearchaeales* median 739 kbp; *Maxwellarchaeales* median 1,030 kbp.

No order encodes a functional electron transport chain — the set of membrane-bound protein complexes (Complex I through IV plus F-type ATP synthase) that generates ATP via oxidative phosphorylation. We examined each complex separately, validating absences against custom archaea-trained HMMs to rule out KEGG-family under-detection of divergent archaeal orthologs (Methods).

*Complex I (NADH:quinone oxidoreductase).* No genome encodes a complete Complex I: nuoA, nuoJ, and nuoK are absent from all 446 genomes even against archaea-trained HMMs, and the remaining subunits are either absent or present at scattered low-confidence levels. nuoF (K00335) is the one exception, recovered in 148 of our 256 *Nanobdellota* genomes with scores up to 789 bits against a custom HMM trained on 200 non-*Nanobdellota* archaeal nuoF sequences — consistent with a retained single-protein role rather than functional NADH:quinone oxidoreductase.

*Complex II (succinate dehydrogenase).* sdhA (K00239) is recovered in 31 of our 256 *Nanobdellota* genomes with scores up to 478 bits against an archaea-trained HMM, but sdhBCD are absent from all 446 genomes. sdhA alone cannot form a functional Complex II and has likely been repurposed, as documented in some anaerobes.

*Complex III (cytochrome bc1).* Absent from archaea as a whole, not a *Nanobdellota*-specific loss: in the 8,776 non-*Nanobdellota* R232 archaeal proteomes we surveyed, petA and petC recruited zero strict hits and petB recruited one.

*Complex IV (cytochrome c oxidase) and F-type ATP synthase.* coxABC and atpABCDEF are absent from all 446 *Nanobdellota* at strict KofamScan threshold. The custom archaea-trained HMM hits observed for K02111 atpA and K02112 atpD (122 *Nanobdellota* each) were verified to be 100% V/A-type ATPase cross-detections of the same proteins identified by K02117 and K02118, reflecting the shared AAA+ ATPase domain rather than a genuine F-type signal.

V/A-type ATPase, present in a subset of *Woesearchaeales* and *Parvarchaeales*, is the only proton-motive machinery in the phylum (Table 1).

Where KofamScan results were flagged as borderline, we validated with archaea-trained custom HMMs (trained on hits from the 8,776 non-*Nanobdellota* HQ R232 archaeal proteomes and searched back against our 256 *Nanobdellota*). The KofamScan crosscheck against UniRef90 validated the core metabolic findings with no major contradictions. Of six KOs previously flagged as “unresolved” at the UniRef90 stage (sdhA, bioB, bioF, hisE, murB, argAB), the custom archaea-trained HMM analysis confirms sdhA as genuinely present (see above); the remaining five have not been reassessed.

### Form III rbcL transfer from archaea to CPR

To assess whether rbcL in our *Nanobdellota* genomes shows any local signal of recent lateral exchange with co-sampled bacteria, we extracted rbcL from the *Patescibacteriota* — the Candidate Phyla Radiation (CPR), a bacterial superphylum whose lineages, like DPANN, are mostly small-genome and host-associated; *Patescibacteriota* is the GTDB phylum name and “CPR” is the original informal name, and the two terms refer to the same group — assembled in the same six samples. Of 632 high-quality (≥90% complete, <5% contamination) single-contig *Patescibacteriota* from our samples, 42 genomes (6.6%) carry at least one strict-threshold K01601 hit.

At mmseqs2 easy-cluster sweep thresholds of 90% and 85% identity (80% mutual coverage), no cluster contains both a *Nanobdellota* rbcL and a *Patescibacteriota* rbcL sequence — recent transfer at high sequence identity is not supported. At 80% identity, a single seven-member cluster (pairwise 80.0–84.1% identity, 95%+ mutual coverage) contains the rbcL of one Baltic Sea water column *Minisyncoccia* genome (B10_u4232805, 99.4% complete, 663 kbp single contig) together with rbcL from six *Nanobdellota* genomes: five *Pacearchaeales* (GW2011-AR1 family) from Baltic and SA1420A samples plus one Baltic *Woesearchaeales*.

Outgroup-rooted phylogeny — using a Form IB plant rbcL set as the outgroup, where Form I is the canonical Calvin-cycle RuBisCO of plants and most cyanobacteria and provides a deep, consistently monophyletic outgroup to the Form III tree we are inferring (Methods *rbcL phylogeny*) — places the *Minisyncoccia* rbcL nine ancestral nodes deep inside an archaeal-majority subclade whose only non-archaeal content is CPR rbcL (no Form III bacteria, no non-*Nanobdellota* archaea, no KEGG references in the subclade), consistent with a single archaea-to-CPR transfer to *Minisyncoccia* from a *Pacearchaeales* or *Woesearchaeales* lineage in our sampled environments.

At the locus level, *Pacearchaeales* SA1420A_u50695000 carries a REP-associated tyrosine transposase (KEGG K07491) four genes downstream of rbcL, while the *Minisyncoccia* rbcL neighborhood is sparse and transposase-free — a pattern compatible with past mobile-element activity at the archaeal locus.

The *Minisyncoccia* (proposed by Nakajima et al. 2025 as a class of the new phylum *Minisyncoccota*; retained here in GTDB R226 *Patescibacteriota* for consistency with our analysis) is the CPR-equivalent class that contains the recently cultured archaea-parasitizing isolate *Minisyncoccus archaeiphilus* (Nakajima et al. 2025), establishing the existence of cultured CPR–archaea parasitism in the lineage. The cultured isolate parasitises a methanogen (*Methanospirillum hungatei*), not a *Nanobdellota*; the relationship reported here is at the class level rather than at the species level, and provides no direct evidence about whether the rbcL-sharing Baltic *Minisyncoccia* parasitises any archaeon. This result does not speak to older transfers or to CPR–archaea exchange outside our sampled environments.

A 4,262-tip rbcL gene tree (built from K01601 hits across our 256 internally-assembled *Nanobdellota* plus 238 high-quality NCBI *Nanobdellota* MAGs (494 genomes total in the *our_nano* pool), our 632 co-sampled *Patescibacteriota*, 1,031 R232 high-quality *Nanobdellota*, 8,776 R232 high-quality non-*Nanobdellota* archaea, 11,567 R232 high-quality *Patescibacteriota*, 317 R232 high-quality Form III bacterial genomes, and 1,498 KEGG K01601 references; LG+R10 model, 296 BMGE-trimmed amino-acid positions, IQ-TREE3) was pruned (Strategy A: dereplicate paralogs; Strategy B: drop target-free deep clades, keeping 5 anchors per major form) to a 1,154-tip subtree, then rooted on the four-tip Form IB plant outgroup and trimmed of those plants to give the 1,150-tip analysis tree on which event enumeration is performed; the 1,150-tip analysis tree retains 391 *Nanobdellota* tips and 320 *Patescibacteriota* tips along with non-*Nanobdellota* archaeal context (Methods *rbcL phylogeny*). Systematic examination of this analysis tree identifies the *Minisyncoccia* case described above as one of nine distinct deep archaea-to-CPR nesting events visible across the 320 *Patescibacteriota* rbcL tips (Table 3). Each event has a CPR tip or small CPR cluster nested inside a ≤21-tip archaeal-majority clade (≥3:1 archaea:CPR under Form IB plant outgroup rooting), comprising twelve CPR tips in total (∼4% of the *Patescibacteriota* rbcL set). Donor archaeal lineages are heterogeneous: four events nest within *Nanobdellota*/DPANN subclades, four nest among non-*Nanobdellota* archaea, and one (Event 9) sits in a mixed *Nanobdellota* + non-*Nanobdellota* archaeal clade (Table 3, Donor lineage column), indicating that archaea-to-CPR rbcL exchange is not restricted to one archaeal donor lineage. A second event in our data involves a *Patescibacteriota* from KR0015B (KR0015B_u24324275), nested within a nine-tip non-*Nanobdellota* archaeal clade, a different archaeal donor lineage than the *Minisyncoccia* case. Per-event UFBoot support at each event’s defining archaea-majority ancestor ranges from 6 to 99 (median 81; Table 3): five events are at UFBoot ≥81, three at 53–59, and one at UFBoot 6 (this latter event is retained for directional tabulation but not interpretable as an individual transfer). Events 5 and 6 share an enclosing ancestor and may represent two related transfers into the same archaeal subclade rather than two fully independent events. The overall pattern — episodic, rare, and lineage-heterogeneous archaea-to-CPR rbcL transfer — is robust to rooting choice; individual event boundaries are subject to the rbcL gene tree’s overall UFBoot split correlation of 0.922 in addition to the per-event support reported in Table 3.

**Table 2.** Putative split tRNA patterns by order across all 208 complete genomes. Rows: amino acids with order-specific missing tRNAs above the tRNAscan-SE archaeal score cutoff (Trp; Val). Values: number of genomes missing tRNA / total genomes per order (and percentage). The remaining standard amino acids are detected in all 208 genomes (after dropping the iMet, Sup, Undet, and SeC categories from the tRNAscan-SE output and retaining pseudogenes). Apparent absences may reflect either genuine loss or below-threshold detection of split (trans-spliced) tRNAs known from *N. equitans* (Randau et al. 2005, *Nature*; canonical set established in Randau, Pearson and Söll 2005, *FEBS Letters*); the canonical *N. equitans* split-locus set comprises Glu, His, Lys, and iMet — neither Trp nor Val is among them, so the patterns reported here are not directly precedented at the locus level.

| Amino<br>acid | Woesearchaeales<br>(123) | Pacearchaeales<br>(54) | Maxwellarchaeales<br>(31) |
| --- | --- | --- | --- |
| Trp | 30/123 (24.4%) | 9/54 (16.7%) | 1/31 (3.2%) |
| Val | 0/123 (0.0%) | 15/54 (27.8%) | 0/31 (0.0%) |

**Table 3.**
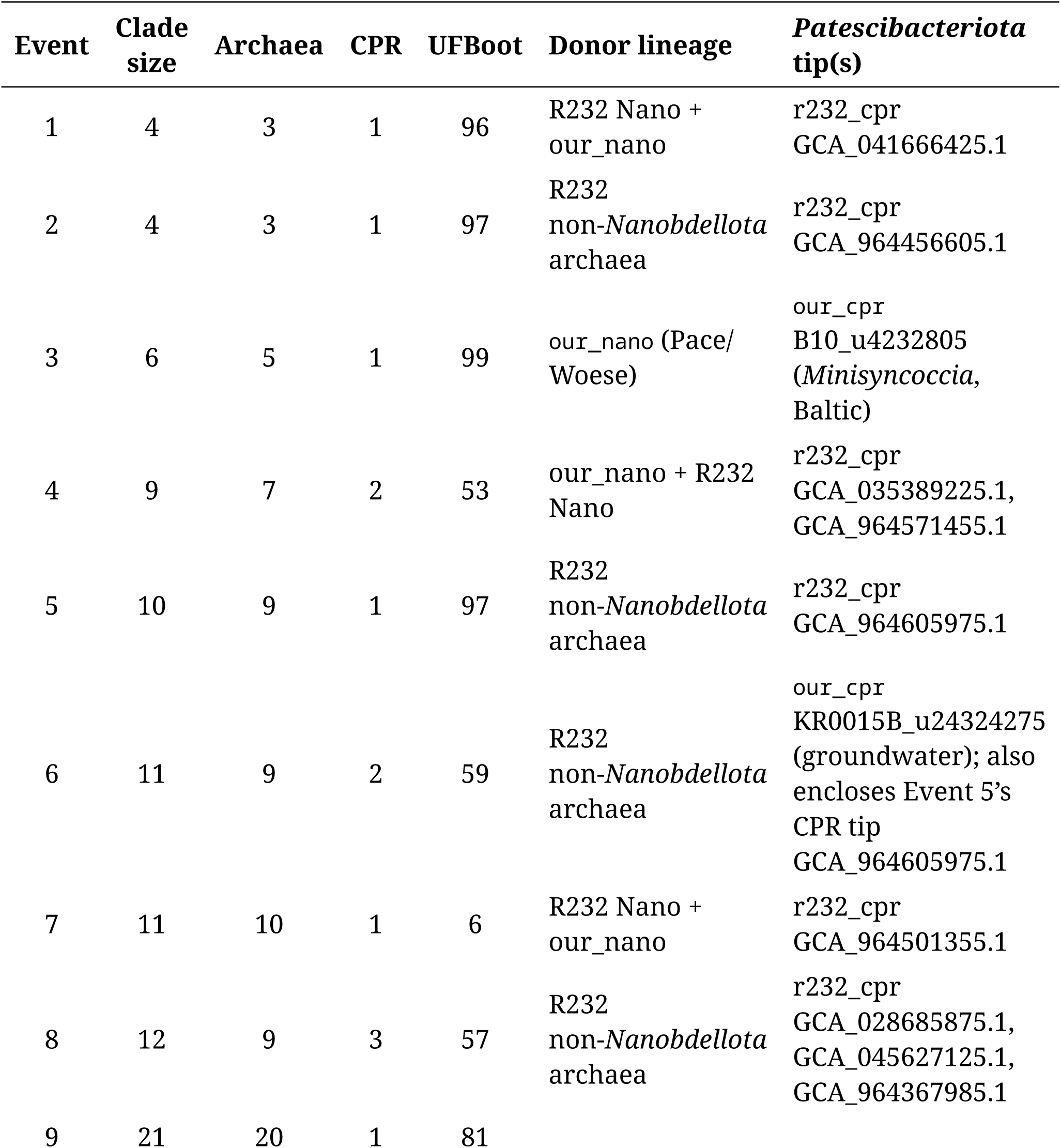
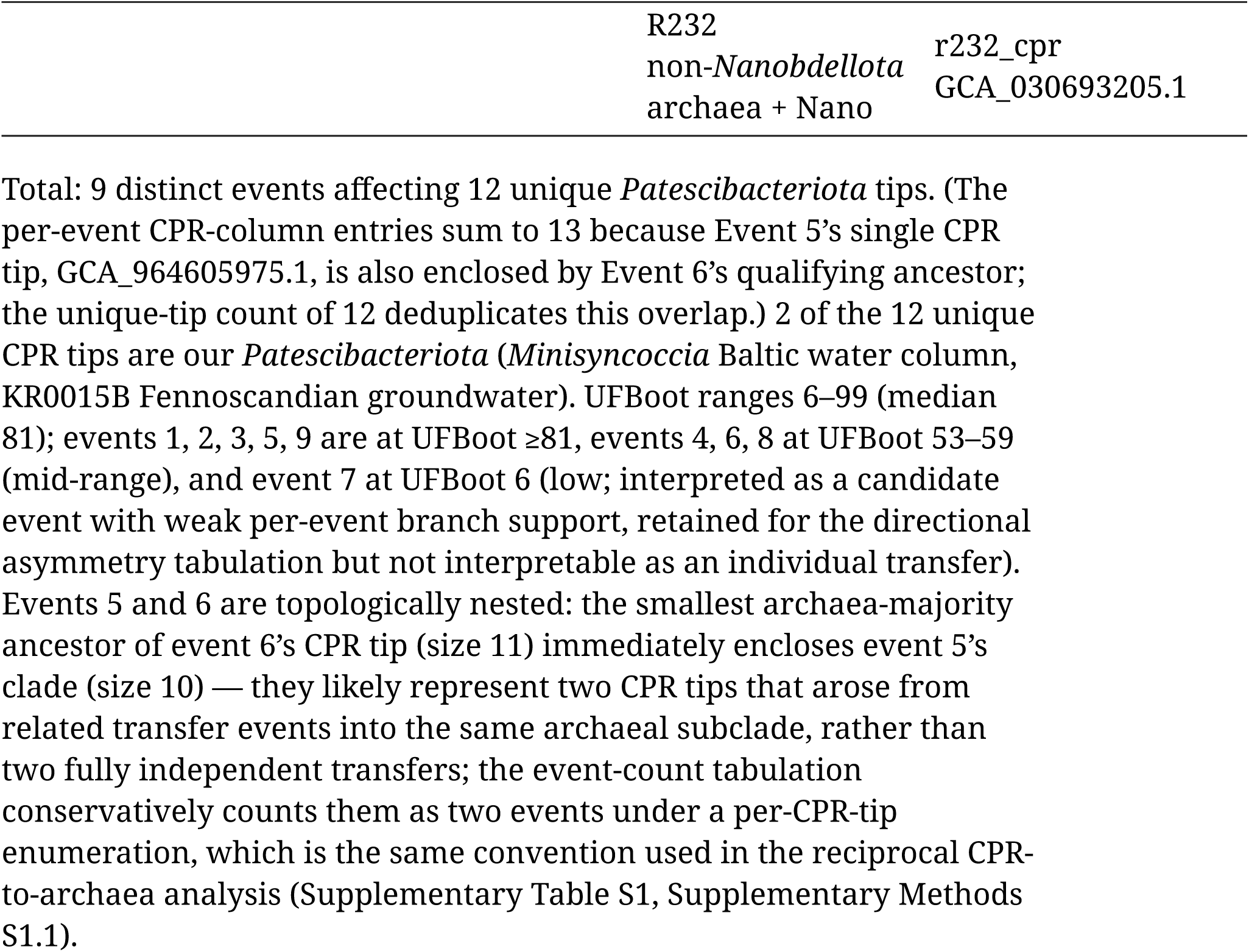
Deep archaea-to-CPR rbcL nesting events visible in the pruned, plant-rooted rbcL gene tree (1,154 tips post-Strategy-A/B reduction to 1,150 leaves after the 4-tip Form IB plant outgroup is removed post-rooting; event enumeration is performed on the 1,150-leaf analysis tree). Each row is one distinct event — a CPR tip or small CPR cluster nested inside a ≤21-tip archaeal-majority clade (≥3:1 archaea:CPR; “qualifying” = clade size 4–21 tips, see Supplementary Methods S1.1). Archaea count includes our *Nanobdellota*, R232 HQ *Nanobdellota*, and R232 HQ non-*Nanobdellota* archaea. “Our” indicates *Patescibacteriota* from the six samples reported here. Donor-lineage abbreviations: *our_nano* = *Nanobdellota* from this study (the six samples); *R232 Nano* = high-quality *Nanobdellota* MAGs from GTDB R232; *R232 non-*Nanobdellota* archaea* = high-quality archaeal MAGs in GTDB R232 from any phylum other than *Nanobdellota*; *Pace/Woese* = a within-*Nanobdellota Pacearchaeales*–*Woesearchaeales* subclade. The UFBoot column is the IQ-TREE3 ultrafast-bootstrap support value at the smallest qualifying archaea-majority ancestor of each event’s CPR tip(s), extracted from the rbcL contree (rubisco_tree_v2/ event_ufboot.py).

To confirm that archaeal *Nanobdellota* Form III rbcL and *Patescibacteriota* Form III rbcL are the same protein family at the sequence level (rather than convergent Form-III-like folds arising from independent evolution), we built two custom HMMs on non-overlapping training sets: one from the 133 strict-K01601 rbcL sequences in our 256 *Nanobdellota* genomes, and one from the 42 strict-K01601 rbcL sequences in our 632 *Patescibacteriota*. Each HMM was then cross-tested against the opposite kingdom’s proteomes. The *Nanobdellota*-trained HMM detected every strict-K01601 *Patescibacteriota* rbcL (42/42 at e<10⁻⁵, against our 632 co-sampled *Patescibacteriota* proteomes) plus 6 additional sub-K01601 hits, for a total of 48 *Patescibacteriota* rbcL detected. The *Patescibacteriota*-trained HMM detected every strict-K01601 *Nanobdellota* rbcL (133/133 at e<10⁻⁵, against our 256 *Nanobdellota* proteomes) plus 15 additional sub-K01601 hits, including one 44-aa pseudogene fragment, for a total of 148 *Nanobdellota* rbcL detected. Own-kingdom bitscores were only 15–20% higher than cross-kingdom bitscores, a profile consistent with a shared ancestral protein family rather than convergent Form-III-like folds. We note that the HMMs were trained on K01601-positive sequences from each kingdom, so this cross-detection test is not strictly orthogonal to the K01601 source filter and is treated as a sanity check rather than a primary result; the same shared-family conclusion is supported directly by the K01601 hits in both kingdoms reported above.

### A single-copy information-processing core

We identified single-copy orthogroups (an orthogroup is a cluster of putatively orthologous proteins across the input genomes; “single-copy” means present at exactly one copy in most or all genomes, the gene set ideal for phylogenomic supermatrix construction) across 206 of the 208 complete genomes — the cohort current at OrthoFinder run time — using OrthoFinder v3.1.4 (Emms and Kelly 2019) with all-vs-all DIAMOND search and MCL clustering at inflation 1.2. The 2 promoted-back *DEMOTED_no_ORC1* genomes were added downstream via hmmsearch under the same calibrated GA threshold used for the NCBI MAGs (Methods *Marker set selection and HMM construction*).

OrthoFinder’s automatic species-tree-orthogroup selection algorithm (`DetermineOrthogroupsForSpeciesTree`; walks a single-copy fraction threshold from 1.0 downward, applying a probabilistic test for hidden paralogy at each step, and stops when at least 100 orthogroups qualify and further relaxation no longer doubles the count per relative threshold-decrease) identified 101 single-copy orthogroups for the concatenated species-tree alignment. All 101 are information-processing or cell-machinery genes: ribosomal proteins of both the small and large subunits, translation initiation and elongation factors, aminoacyl-tRNA synthetases, a subset of RNA polymerase subunits (RpoA paralogs preclude single-copy classification of the largest two subunits; see Methods *ar71 phylogeny*), RNA processing enzymes, replication proteins (ORC1/Cdc6, DNA primase DnaG, replication factor C, flap endonuclease), cell division proteins, a proteasome subunit, and the preprotein translocase SecY. No gene of central or energy metabolism — glycolysis, TCA cycle, amino acid biosynthesis, electron transport, or RuBisCO — is conserved as single-copy across the phylum. Combined with the order-level metabolic divergence documented above, the minimal *Nanobdellota* cell is defined by replication, transcription, translation, and cell division rather than by any conserved metabolic framework.

The *Nanobdellota* core is exceptionally divergent at the sequence level. Median within-phylum pairwise amino acid identity, sampled across 500 random pairs per gene on trimmed alignments, ranges from 31.4% (ribosomal protein eS24) to 65.9% (ribosomal protein S11), with an overall median of 47.4% and individual minimum pairs as low as 13.2% (OG0000296, structurally a winged-helix–turn–helix transcriptional regulator by Foldseek vs AFDB; revised from a sequence-based VapC9 PIN-like call). BMGE v2.0 (Criscuolo and Gribaldo 2010) trimming with the BLOSUM30 substitution matrix retained only 42.8% of alignment positions across the 66-OG quality-filtered core (26,516 raw to 11,352 trimmed amino acid positions), compared to 60–70% typical for bacterial supermatrices at comparable taxonomic scope. This level of within-phylum divergence reflects the fast-evolving, reduced-genome character of DPANN archaea and motivates the construction of phylum-specific phylogenetic markers rather than reliance on the broadly-trained all-Archaea marker sets ar53 and ar122 used by GTDB.

### A Nanobdellota-specific phylogenomic marker set

The *Nanobdellota* marker set (ar71) was built from the 101 single-copy orthogroups by an iterative HMM-refinement procedure tuned to the within-*Nanobdellota* single-copy criterion. For each orthogroup, an initial profile HMM was built from the 206-genome single-copy alignment with MAFFT v7.525 L-INS-i and HMMER v3.4 (Eddy 2011) `hmmbuild`, then expanded by aligning high-scoring hits from a 1,288-genome *Nanobdellota* calibration pool (our complete-genome set together with all high-quality *Nanobdellota* MAGs in GTDB R232) into the alignment with MAFFT `--add --keeplength` and rebuilt; iteration continued until the single-copy hit set converged or eight rounds were reached, with each marker rolled back to the iteration that maximized single-copy recovery on the 1,288-genome pool. Hits were accepted only at envelope coverage ≥70% of HMM length, dropping partial / fragment matches that inflate apparent paralogy. Per-marker GA/NC bit-score thresholds were calibrated against the 206-genome OrthoFinder OG-membership ground truth (TPs = OG members; FPs = all other 206-genome proteins): GA = midpoint of min(TP) and max(FP) when separable, else 1st percentile of TPs; NC = max(FP). Markers were retained only if the calibrated HMM achieved single-copy ≥1,100/1,288, multi-copy rate ≤5%, and GA ≥30 on the 1,288-genome pool — a quality filter that selects for markers that generalise beyond the 206-genome training set across all named *Nanobdellota* orders. 71 of the 101 orthogroups passed this filter; the 30 that did not either failed the multi-copy criterion (paralog families distributed across subsets of *Nanobdellota*), the recovery threshold (under-detection in distantly-related orders), or the score-separability criterion (no clean GA boundary between true positives and false positives). The final 71-marker set is the ar71 marker set used throughout this paper. The marker HMMs with embedded GA/NC thresholds are deposited at `custom_hmms/ar71/`.

Comparison to the standard archaeal marker sets used by GTDB reveals both substantial overlap and genuine novelty. Of the 71 ar71 markers, roughly half overlap with the ar53 set (53 archaeal markers, used by GTDB since R07-RS207) and the majority overlap with ar122 (the earlier 122-marker set); the remainder are markers identified by our within-*Nanobdellota* calibration that are not part of either GTDB set. The novel markers were retained because they passed the within-*Nanobdellota* single-copy and score-separability criteria across all named *Nanobdellota* orders represented in the 1,288-genome calibration pool. They comprise an S1 RNA-binding motif, a GNAT-fold N-acetyltransferase, a proteasome endopeptidase, the ribosome biogenesis factor KRR1, phenylalanyl-tRNA synthetase α-subunit, a winged-helix–turn–helix transcriptional regulator, a protein-synthesizing GTPase, and a small number of uncharacterized proteins; OrthoFinder OG identifiers are listed in `custom_hmms/ar71/index.tsv`.

Two of these architectures — the GNAT-fold N-acetyltransferase (OG0000275) and the winged-HTH transcriptional regulator (OG0000296) — were assigned by Foldseek structural search of the OG-representative ESMFold model against the AlphaFold Database (top-5-consensus annotation; query PDBs and result TSV deposited at foldseek_dark_ogs/), revising sequence-based annotations of *PRC-barrel domain* and *VapC9 PIN-like nuclease* respectively; the remaining labels are sequence-based.

To characterise the 71-marker set at the 446-taxon scale, per-marker single-copy amino acid sequences were extracted from all 446 taxa (208 complete genomes — 206 via OrthoFinder orthogroup membership and 2 via hmmsearch under the calibrated GA threshold; 238 NCBI MAGs via hmmsearch with the same calibrated GA threshold), aligned with MAFFT v7.525 L-INS-i, and trimmed with BMGE v2.0 under BLOSUM30. Minimum per-genome occupancy across the 446-taxon set was 54 of 71 markers (76%); all 446 taxa were retained for downstream characterization. The main phylogeny used in this paper is the 71-marker ar71 tree on 1,239 R232-extended taxa described in Methods *ar71 phylogeny*.

### tRNA, rRNA, and assembly frameshifts

Infernal cmsearch detected a single 16S–23S–5S rRNA operon in 203 of the 208 complete genomes; the three exceptions (B10_u24296491, B5_u20127012, SA1420A_u28577139) carry SSU and LSU but no detectable 5S above the cmsearch inclusion threshold. One *Woesearchaeales* genome (B10_u28622061) carries three SSU, three LSU, and six 5S rRNA loci; the underlying 16S–23S–5S operonic arrangement is not resolvable from cmsearch coordinates alone. tRNAscan-SE identified 26–54 tRNAs per genome (median 35); 96 of 208 genomes encode tRNAs for all 20 standard amino acids. The remaining genomes show order-level missing-tRNA patterns (tRNA-Trp missing from 30/123 *Woesearchaeales*, 9/54 *Pacearchaeales*, and 1/31 *Maxwellarchaeales*; tRNA-Val missing from 15/54 *Pacearchaeales* only) (Table 2). Independently evolved split (trans-spliced) tRNAs are a documented phenomenon in DPANN: Randau et al. (2005, *Nature*) initially identified split-half-encoded tRNAs in *N. equitans* by computational prediction (Glu, His, Trp, iMet), with the cognate tRNA reconstituted from independently encoded 5’- and 3’-halves below the threshold of standard single-locus tRNA detection; Randau, Pearson and Söll (2005, *FEBS Letters*) subsequently established the biochemically verified canonical set as tRNA-Glu, tRNA-His, tRNA-Lys, and initiator tRNA-Met; the originally predicted tRNA-Trp half was corrected to tRNA-Lys (the intron-containing tRNA-Trp gene was misidentified as a split half in the initial annotation). Neither Trp nor Val is among the canonical four amino acids for which an *N. equitans* split-locus arrangement is established; the Trp-and Val-missing patterns reported here are therefore consistent either with order-level partial-isoacceptor loss, a paralogous-locus detection failure, or a *Nanobdellota*-specific extension of the same split-tRNA mechanism to additional loci — these alternatives cannot be discriminated from our data alone.

As a quality assessment of the Oxford Nanopore assemblies (Watson and Warr 2019), assembly frameshifts totalled 1,214 across the 208 complete genomes (325 split-gene events, adjacent same-strand ORFs sharing a UniRef90 best-hit target separated by ≤200 bp; 889 truncated-with-gap events, a UniRef90-truncated gene followed by a same-strand intergenic gap of plausible size; Methods *Frameshift detection*). The mean per-genome rate was 5.9 events (5.47 across the 123 Baltic complete genomes vs 6.52 across the 85 Fennoscandian groundwater complete genomes, a 19% higher rate in groundwater that is consistent with the ∼2× lower mean myloasm Depth1 (≥99% identity) of the groundwater Nano contigs at 60.6× vs 33.5×; KR0015B 12.6× alone; though a sequencing-coverage vs assembly-quality vs biology decomposition cannot be made from these data); 3 of 208 genomes carried no detected frameshift, representing ∼0.4% of predicted loci.

Some fraction of these events represent genuine biology (programmed frameshifts in genes such as *prfB*, recent pseudogenization along the host-association trajectory, and rotation-seam artifacts at the wrap point) rather than sequencing error, and the count is interpreted as an upper bound on assembly-derived frameshifts rather than a strict assembly-error rate.

## Discussion

### Complete genomes transform a data-poor phylum

Prior to this work, the phylum *Nanobdellota* was represented by four complete genomes (all from thermophilic environments) along with one additional single-contig high-quality MAG whose closure status and replication-origin placement were not established. The 208 complete genomes presented here represent a 52-fold increase in genuinely complete *Nanobdellota* genomes and, critically, provide the first complete environmental genomes for the three orders that dominate aquatic and subsurface habitats. Completeness is not merely a quality metric for these organisms. At genome sizes of 553 kbp to 2,676 kbp with ∼90% coding density, the distinction between a genuine gene absence and an assembly gap is load-bearing for biological interpretation. The metabolic absences we report — zero electron transport chain, zero citrate synthase, zero F-type ATPase — are interpretively strong precisely because they come from complete genomes.

### A minimal Nanobdellota cell defined by its information-processing core

The 101-orthogroup information-processing core, derived from our 208 complete genomes and extended to the high-quality *Nanobdellota* MAGs in GTDB R232 via phylum-specific HMMs, defines the universally conserved functional core of the *Nanobdellota* cell. The core is exclusively information-processing and cell-machinery (ribosomal proteins, translation factors, aminoacyl-tRNA synthetases, RNA polymerase subunits, replication and cell-division proteins, a proteasome subunit, the SecY translocase); no gene of central or energy metabolism is conserved as single-copy across the phylum. The functional silhouette of a phylum-wide minimal *Nanobdellota* cell is therefore: replicate, transcribe, translate, and divide; everything else is variable. This silhouette is consistent at the extreme end with Harada et al.’s description of *Candidatus Sukunaarchaeum* mirabile, a 238-kbp archaeon retaining primarily replicative machinery (phylogenetic placement uncertain; Harada et al. 2025, preprint). We have not performed an OG-by-OG overlap with the *Sukunaarchaeum* annotation in the present analysis. The silhouette helps frame the order-level metabolic divergence below: orders differ in what they retain *on top of* this shared core, not in the identity of the core itself.

### Divergent metabolic strategies across orders

The complete-genome dataset resolves a fundamental metabolic divergence between orders within a single phylum, observed independently in both Baltic Sea water column and Fennoscandian groundwater. *Woesearchaeales* encodes the broadest metabolic toolkit within the phylum: partial glycolysis in most genomes, V/A-type ATPase in approximately one-third, partial amino acid biosynthesis and cofactor biosynthesis in subsets, and glycogen storage. Even so, the absence of the electron transport chain means *Woesearchaeales* cannot conduct oxidative phosphorylation. *Pacearchaeales* and *Maxwellarchaeales*, by contrast, retain a much narrower toolkit dominated by RuBisCO, PEP synthase, and ferredoxin on top of the information-processing core, with scattered remnants of glycolysis (in *Pacearchaeales*) and partial pyruvate:ferredoxin oxidoreductase plus succinyl-CoA synthetase (in subsets of *Maxwellarchaeales*); no order encodes a complete autonomous energy-metabolism pathway. The three orders clearly represent divergent adaptations to the same fundamental constraint — survival with minimal genomes and no electron transport chain — using different metabolic strategies. The most distinctive order-level feature of this divergence is rbcL distribution: universal in *Pacearchaeales* (100%), frequent in *Maxwellarchaeales* (86%), but rare in the larger-genome *Woesearchaeales* (11%). This pattern is not parsimonious under a simple gene-loss-with-genome-shrinkage scenario, and is examined in detail in the RuBisCO section below.

This divide has implications for host dependency. *Pacearchaeales* and *Maxwellarchaeales* lack ATPase and encode at most scattered glycolytic enzymes (pyruvate kinase in 9/54 *Pacearchaeales* and 6/31 *Maxwellarchaeales*), suggesting a high degree of host dependency for energy; the minority retention of pyruvate kinase is not by itself a route to autonomous substrate-level ATP, since the PEP substrate is not generated by upstream glycolysis in these orders (see the energy-budget analysis below). *Woesearchaeales* retains a broader gene complement for energy and substrate metabolism in a subset of genomes; whether this implies a less obligate host relationship cannot be inferred from gene content alone, since closely-related cultured DPANN often retain machinery that is functional only in the presence of a host. *Candidatus Sukunaarchaeum* mirabile (introduced above; Harada et al. 2025, preprint) sits at the extreme end of this host-dependency gradient at 238 kbp; its OG-level overlap with the 101-OG core defined here is left as a future-work item.

The within-phylum metabolic gradient documented here is internal to *Nanobdellota* and does not directly test the inter-phylum DPANN ancestry question addressed by Baker et al. (2025), since rooting at the phylum stem requires a deeper outgroup than our supermatrix supplies. Within *Nanobdellota*, the metabolic capacity required for host-independent life is partially retained in *Woesearchaeales* (glycolysis and V/A-type ATPase in subsets of genomes) but essentially absent in *Pacearchaeales* and *Maxwellarchaeales*. The metabolic absences shared across all 446 genomes (no electron transport chain, no F-type ATPase, no citrate synthase, regardless of order) establish that order-level metabolic differences are differences in retained machinery against a phylum-wide baseline of loss. The within-phylum gradient is consistent with at least two scenarios that the present data do not distinguish: (i) a reductive trajectory in which *Pacearchaeales* and *Maxwellarchaeales* have lost more capabilities than *Woesearchaeales*, the within-phylum complement of the reductive-from-free-living account favoured by Baker et al. (2025); or (ii) a scenario in which *Pacearchaeales* and *Maxwellarchaeales* retain a closer-to-ancestral minimal-genome state and *Woesearchaeales* lineages retain or have re-acquired capabilities lost in their sister orders. We do not assert a direction of evolutionary change at the phylum stem (our supermatrix trees are midpoint-rooted; rooting on a deeper outgroup would be required to claim that the phylum-wide absences predate order-level diversification rather than the converse), and the within-phylum gradient stands as an observation that any larger-scale account of DPANN evolution must accommodate, regardless of which scenario for the phylum stem eventually proves correct.

The complete absence of aerobic respiration across the 208 complete genomes is a phylum-level trait, not an order-specific loss; in the 238 NCBI MAGs, the absences are also consistent across orders, but MAG incompleteness means the strength of the negative claim is set by the 208 complete genomes alone. Energy generation inferred from the annotated gene content is limited to substrate-level phosphorylation and host-derived ATP; novel pathways encoded by the uncharacterized fraction of these proteomes cannot be ruled out.

Considered as an energy-budget question, the *Pacearchaeales* gene complement (54 complete genomes from this study) does not support self-sustained ATP generation by canonical pathways. F-type ATP synthase and V/A-type ATPase are absent (0/54 in both); pyruvate:ferredoxin oxidoreductase is incomplete in every genome (3/54 with α + β + δ subunits at strict KofamScan threshold but γ subunit absent throughout, so no genome encodes the canonical four-subunit assembly); and pyruvate kinase (9/54) provides the only detectable substrate-level phosphorylation enzyme, but acts on a phosphoenolpyruvate substrate that no *Pacearchaeales* genome can generate from glycolysis (GAPDH 13/54, phosphoglycerate kinase 13/54, with no genome encoding a majority of the upstream pathway).

*Maxwellarchaeales* (31 complete genomes) retains more of the same machinery in a subset of members: 28/31 PEP synthase, 17/13/13/11 of 31 for the four PFOR subunits (α/β/δ/γ respectively), 6/31 pyruvate kinase, and 6/31 succinyl-CoA synthetase α. This admits partial substrate-level and PFOR/PEP-synthase-coupled pathways in subsets of genomes, with the in vivo direction (forward / reverse) of PEP synthase and PFOR not characterized by gene content alone. In numerical summary, 11 of 31 *Maxwellarchaeales* encode the canonical four-subunit PFOR versus 0 of 54 *Pacearchaeales* — the sharpest single-pathway difference between the two orders. Across both small-genome orders, the most parsimonious lifestyle reading remains obligate host-coupled, consistent with the *N. equitans* / *N. aerobiophila* ectosymbiont mode (Huber et al. 2002; Kato et al. 2022) or the nanotube-mediated transfer mode documented in cryo-EM (Johnson et al. 2024), rather than a free-living syntrophic mode; the latter would require sustained autonomous ATP generation that the gene content does not support.

Three caveats apply to the metabolic findings. First, the 500 kbp size filter combined with the CheckM2 high-quality thresholds could in principle exclude small *Nanobdellota* genomes (*N. equitans* is 491 kbp); however, CheckM2 assessment of all 58 *Nanobdellota* contigs in the 400–500 kbp range from Baltic samples found none that were high-quality, and skani confirmed that all were fragments of species already represented in the complete-genome collection. Second, the NCBI genomes are fragmented MAGs, and metabolic absences in the NCBI-only orders should be treated with appropriate caution. Third, our 208 complete genomes derive from two regions — the Baltic Sea water column (four samples) and Fennoscandian groundwater at 69 and 201 m below sea level (two Äspö samples) — so claims about the metabolic state of *Pacearchaeales* and *Maxwellarchaeales* or their host-dependency landscapes pertain to lineages sampled here and should be tested as additional regions are surveyed at complete-genome resolution.

### Form III RuBisCO: function, distribution, and origin

The Form III RuBisCO in *Nanobdellota* (rbcL without rbcS or PRK) is most parsimoniously interpreted as nucleotide salvage by analogy with the AMP-recycling pathway characterized in *Thermococcus kodakarensis* (Sato et al. 2007), where RuBisCO catalyzes the carboxylation of ribulose-1,5-bisphosphate derived from AMP degradation to yield 3-phosphoglycerate. The absence of rbcS (verified by both KofamScan and Foldseek vs AlphaFold-DB rbcS references) together with the absence of any KofamScan ortholog call for PRK across all 446 genomes rules out the Calvin cycle. The full *T. kodakarensis* AMP-shunt is not strictly transposable to the *Pacearchaeales* / *Maxwellarchaeales* metabolic background — *T. kodakarensis* retains a complete glycolytic / gluconeogenic apparatus that consumes the 3-phosphoglycerate output, whereas glycolysis is largely absent in these orders — so the downstream fate of the 3-PGA produced in our genomes is unresolved. Form III RuBisCO is consistent with a salvage role in these organisms, but the precise downstream metabolic context awaits direct biochemical characterization in a host-associated DPANN background.

The order-specific prevalence — *Pacearchaeales* 100% in complete genomes (89% combined with NCBI), *Maxwellarchaeales* 86% (81% combined), *Woesearchaeales* 11% (11% combined), confirmed across Baltic Sea, Fennoscandian groundwater, and NCBI genomes — is the first demonstration of this pattern at phylum scale with complete genomes. The anti-correlation between RuBisCO prevalence and glycolytic capacity (r =-0.577) is descriptive of the between-order pattern; because the correlation is structured by order-level phylogenetic divergence, a within-lineage functional trade-off cannot be inferred from these data alone.

This distribution is not parsimonious under a simple gene-loss-with-genome-shrinkage scenario: if rbcL were lost progressively with decreasing genome size, the largest-genome order (*Woesearchaeales*, median 1,409 kbp) should retain it at the highest rate, not the lowest. Two scenarios can in principle account for the observed pattern: lateral acquisition of rbcL in the *Pacearchaeales* and *Maxwellarchaeales* lineages (as suggested by Jaffe et al. 2019 for DPANN archaea broadly), or ancestral presence of rbcL in the *Nanobdellota* common ancestor with repeated independent loss along the *Woesearchaeales* backbone. Within *Woesearchaeales*, rbcL-carrying genomes form a phylogenetic subcluster — the mean pairwise patristic distance (the tree-path distance summed along branches between two tips) among rbcL-carrying *Woesearchaeales* is approximately 11% shorter than among rbcL-lacking *Woesearchaeales* (`phylogeny/scripts/ rbcl_patristic.py`, computed across all 728 Woese tips of the ar71 tree with strict-K01601-positive vs negative annotation) — indicating that rbcL presence is a coherent lineage-level trait within the order rather than a scattered per-genome event. The gene-tree analysis below refines the interpretation of this pattern.

The rbcL gene tree, rooted on a Form IB plant outgroup, identifies nine candidate archaea-to-CPR transfer events distributed across ∼4% of the *Patescibacteriota* rbcL tips we examined (Table 3); a reciprocal analysis under the same topological criteria identifies only two candidate CPR-to-archaea events at any UFBoot level — both below the conventional UFBoot ≥ 80 confidence threshold (S1.1 at UFBoot 57, S1.2 at UFBoot 41; three R232 HQ *Nanobdellota* rbcL tips nested inside CPR-majority clades of 12–17 tips; Supplementary Table S1) — a qualitative asymmetry between nine archaea-to-CPR candidates and two CPR-to-archaea candidates. This asymmetry sits against a backdrop of 66 distinct pure-CPR subclades (≥10 tips, ≥90% CPR, zero *Nanobdellota*; Supplementary Note S1) in the same tree, indicating that CPR carries extensive vertically-inherited Form III rbcL of its own and that the transfer events we identify are interlopers into an otherwise taxonomically coherent CPR subtree. This partially addresses the question raised by Jaffe et al. (2019) of whether DPANN archaea and CPR bacteria have exchanged rbcL at deeper phylogenetic scales: at the scale of the *Nanobdellota* phylum and co-sampled *Patescibacteriota*, archaea-to-CPR transfer is more frequently identified than the reverse direction in our data.

We do not interpret the 9-vs-2 candidate count as a calibrated directionality estimate. There is a structural sampling asymmetry: at the source-pool level the archaeal Form III rbcL pool (4,268 hits across 10,301 archaeal search-space genomes; 494 `our_nano` + 1,031 R232 Nano + 8,776 R232 non-*Nanobdellota* archaea) is ≈10× larger than the CPR Form III rbcL pool (414 hits across 12,199 CPR search-space genomes; 632 `our_cpr` + 11,567 R232 cpr; Methods *rbcL phylogeny*), and even after Strategy A/B pruning and outgroup trimming to the 1,150-tip plants-trimmed analysis tree the archaea:CPR tip ratio remains ≈2.6:1 (391 Nano + 421 non-Nano archaea + 18 KEGG references: 320 CPR). The candidate counts are also sensitive to the choice of clade-size cap (≤21 tips), majority threshold (≥3:1), and the topological-rooting choice on a tree whose UFBoot split correlation reached 0.922, below IQ-TREE’s`-bcor 0.99` workflow default but still a stable per-branch bootstrap regime. Restricting both directions to UFBoot ≥ 80 yields five forward events (Events 1, 2, 3, 5, 9 at UFBoot 96, 97, 99, 97, 81; five CPR tips) versus zero reverse events (both reverse candidates S1.1 at UFBoot 57 and S1.2 at UFBoot 41 fall below threshold; Supplementary Table S1, UFBoot column). The qualitative asymmetry — multiple archaea-to-CPR candidates at any UFBoot level versus two reverse candidates both below UFBoot 60 — is preserved across the unfiltered and ≥ 80 UFBoot views. A label-randomization permutation test (10,000 permutations of the cpr / nano / arch_other labels among their tip positions, leaving the 18 KEGG and outgroup-residual tips fixed; Methods *rbcL directionality permutation test*) gives null forward and reverse counts of 109 ± 7 and 18 ± 4 (mean ± SD) respectively, with a null mean asymmetry of 90 events that is never below 54 across the 10,000 permutations. The observed 9-vs-2 asymmetry is in the same direction as the null but smaller in magnitude than what the source-pool-size structural asymmetry alone would predict; we therefore report the asymmetry as a directional pattern consistent with archaea-to-CPR being the more frequently identified direction in our data rather than as a calibrated rate, and treat the per-event phylogenetic structure (clade nesting, UFBoot support, donor lineage heterogeneity) as the primary support for the individual events rather than the count comparison. Notably, none of our 208 complete *Nanobdellota* rbcL tips fall inside CPR-majority clades; the two reverse-direction candidates (S1.1, S1.2) are both R232 MAG tips, suggesting that complete-genome resolution on the archaeal side does not surface reverse transfers at the per-clade level we examined.

The sequence-identity and phylogenetic relationship between our *Nanobdellota* rbcL and the rbcL carried by *Patescibacteriota* from the same environments (Results) identifies a specific, bounded signal of inter-kingdom transfer. At high identity thresholds (≥85%) no cross-kingdom clusters form, so recent transfer at high sequence identity is not supported. At 80% identity a single cross-kingdom cluster is observed, involving one Baltic Sea water column *Minisyncoccia* genome and six co-sampled *Nanobdellota*. This places the genomic transfer signal we describe within the CPR-equivalent class (proposed by Nakajima et al. 2025 as *Minisyncoccia* class. nov. of the new phylum *Minisyncoccota*; retained here in GTDB R226 *Patescibacteriota* for consistency) that contains the recently cultured archaea-parasitizing isolate *Minisyncoccus archaeiphilus* (Nakajima et al. 2025). The cultured isolate parasitizes a methanogen and is taxonomically related to our Baltic *Minisyncoccia* at the class level only; it provides a parasitic precedent for the lineage but does not directly speak to the host of the Baltic genome we sampled. Archaeal donor lineages are heterogeneous across the nine events, and individual event boundaries are subject to the rbcL gene tree’s UFBoot correlation (0.922 split correlation, below IQ-TREE’s `-bcor 0.99` workflow default) and to the choice of plant Form-IB outgroup; these are described as candidate events rather than confidence-bounded estimates. Within these caveats, the imbalance is suggestive of an asymmetry in the historical net flux rather than evidence of a strict directional rule. If the asymmetry survives further sampling and tree-resolution improvements, the biological reading consistent with our data, and with the cross-domain parasitic interface documented for the *Minisyncoccia* (Nakajima et al. 2025), is that the archaeal–CPR interface has been a predominantly archaea-to-CPR conduit for Form III rbcL, plausibly along the same parasitic or syntrophic contacts that bring small-genome archaea and bacteria into physical proximity.

The rbcL gene tree resolves this question. Our 391 *Nanobdellota* rbcL tips do not form a monophyletic clade in the analysis tree: their MRCA encompasses the entire 1,150-tip plants-trimmed analysis tree, and the three principal orders each scatter across multiple distinct archaeal subclades interleaved with non-*Nanobdellota* archaeal rbcL. The pattern is consistent with multiple independent lateral acquisitions of rbcL from diverse archaeal donor lineages (Jaffe et al. 2019), rather than ancestral retention of a single ancestrally inherited rbcL with subsequent order-specific loss. The observation reported above that rbcL-carrying *Woesearchaeales* form a phylogenetic subcluster in the species tree (Results) is compatible with two alternatives that the species-tree geometry alone cannot distinguish: (i) a single ancestral rbcL acquisition by the subclade with subsequent vertical inheritance; (ii) repeated rbcL acquisition into the same *Woesearchaeales* subclade from multiple donor lineages. The rbcL gene tree’s scatter of the 391 *Nanobdellota* tips across multiple distinct archaeal subclades — interleaved with non-*Nanobdellota* archaeal rbcL — is what discriminates these alternatives, favouring (ii): a particular *Woesearchaeales* subclade appears to have been a repeated sink for rbcL acquisition from multiple archaeal donors, rather than a lineage that ancestrally carried rbcL.

### Biogeography and phylogenomic placement of orders

The 208 complete genomes from six samples show a sharp biogeographic pattern. The same three orders are present in both Baltic Sea water column and Fennoscandian groundwater at 69–201 m below sea level, with comparable metabolic capabilities in both environments. Yet at the species level, all 145 species are endemic to their environment: 68 Baltic-only, 25 KR0015B-only, 49 SA1420A-only, and 3 species shared between the two Äspö sites (KR0015B and SA1420A); i.e., 74 of 145 species are groundwater-only and split across the two Äspö sites with only 3 cross-site species. Zero species overlap between Baltic Sea and Fennoscandian groundwater, and zero overlap with the 238 NCBI genomes. The closest Baltic–NCBI match is 85.1% ANI, well below the species boundary.

The presence of the same three orders in both environments — despite species-level endemism between the Baltic and Äspö environments — is consistent with the order-level metabolic strategies (glycolysis-encoding *Woesearchaeales*, RuBisCO-bearing *Pacearchaeales* and *Maxwellarchaeales*) having been established before the environmental divergence. The three species shared between KR0015B and SA1420A are consistent with the physical proximity of the two Äspö sites, while the lack of surface–subsurface species sharing suggests limited exchange across that environmental boundary. The only NCBI genomes with any detectable ANI to our genomes are also from Fennoscandian groundwater (Olkiluoto, Finland; Äspö, Sweden), consistent with a shared subsurface Fennoscandian microbiome. Dopson et al. (2024) reported that *Nanobdellota* (as *Nanoarchaeota*) dominated among novel archaeal taxa in Fennoscandian groundwaters from the same facility that produced our KR0015B and SA1420A samples.

The phylum-wide phylogeny separates the three environmental aquatic orders (*Woesearchaeales*, *Pacearchaeales*, *Maxwellarchaeales*), together with UBA10117, into a derived clade. The thermophilic and extreme-environment orders (*Nanobdellales*, *Jingweiarchaeales*, *Parvarchaeales*, *Tiddalikarchaeales*, CSSed11-243R1, WJKC01, JAPDLS01, JAQUSD01) are basal under midpoint rooting (Results *Phylogenomic structure*; Figure 1). Midpoint rooting assumes approximately clock-like evolution, which is a strong assumption for a phylum that spans hyperthermophiles to cold marine water. The basal placement of thermophilic orders should therefore be read as the configuration our data supports under that rooting choice rather than as a confident claim about ancestral state; an outgroup-rooted analysis on a deeper-archaea anchor would be needed to test the *direction* (basal-vs-derived) directly. Robust rooting at this evolutionary depth requires the long-branch-attraction and compositional-bias controls of the kind applied by Baker et al. (2025), which are beyond the scope of the within-phylum analysis we present. Per-order monophyly, by contrast, is a property of the *unrooted bipartitions* in the inferred tree and is therefore robust to the rooting choice: each named multi-tip *Nanobdellota* order, including *Woesearchaeales*, forms a single bipartition in both the ar53 and ar71 trees — the basis for retaining the existing GTDB order-level circumscription documented in *Woesearchaeales is recovered monophyletic* below.

### A phylum-specific phylogenomic marker set for Nanobdellota

The availability of 208 complete *Nanobdellota* genomes enabled the construction of ar71, a phylum-specific phylogenomic marker set that differs in composition from the broadly-trained archaeal marker sets used by GTDB. ar71 was built by an iterative HMM-refinement procedure with calibrated per-marker GA/NC bit-score thresholds and a domain-coverage filter (Methods *Marker set selection and HMM construction*); 71 of 101 candidate single-copy orthogroups passed all three within-*Nanobdellota* criteria (single-copy ≥1,100/1,288 across the calibration pool, multi-copy rate ≤5%, score-separable). The 30 OGs that did not pass were excluded for paralog families distributed across *Nanobdellota* subsets (e.g., RNA polymerase subunits A’ and A’’), under-detection in distantly-related orders, or lack of a clean GA boundary between true positives and false positives. The result is a marker set tuned to within-*Nanobdellota* single-copy resolution at the cost of broader cross-phylum interpretability. These mismatches with GTDB’s broader archaeal sets are not errors in either marker set but reflect an inherent tension between domain-wide and lineage-specific phylogenomic tools: markers conserved across all Archaea must remain interpretable in lineages as divergent as *Thaumarchaeota*, Haloarchaea, and *Nanobdellota*, which selects for a subset of markers that are necessarily dilute within any single phylum.

The markers unique to our set include several that are functionally informative — a winged-helix–turn–helix transcriptional regulator, phenylalanyl-tRNA synthetase α-subunit, a proteasome endopeptidase, a ribosome biogenesis factor (KRR1), and a protein-synthesizing GTPase — alongside a small number of uncharacterized proteins that may represent *Nanobdellota*-specific conserved orthologs of currently unknown function. Within-phylum median pairwise amino acid identity across the calibrated marker alignments is 47.4%, with individual pairs dropping below 15%.

### Woesearchaeales is recovered monophyletic

Whether a named taxon is monophyletic — that is, whether its members form a single clade rather than being interrupted by tips of other named taxa — is the load-bearing constraint under both the SeqCode and the International Code of Nomenclature of Prokaryotes for retaining a taxon at its current rank. We test order-level monophyly in the unrooted bipartitions of our two primary inferences, the ar53 phylogeny (444-tip ML tree from a 446-taxon alignment) and the ar71 phylogeny (1,239 taxa), since the bipartition test is rooting-independent and therefore the appropriate criterion for a topology assertion that must be robust to root placement.

*Woesearchaeales* as currently defined by GTDB is recovered as a single bipartition in both trees: the 205 *Woesearchaeales* tips at the 444-tip ar53 scale and the 728 *Woesearchaeales* tips at the 1,239-tip ar71 scale each form one side of an edge of their respective inferred trees, with all non-*Woesearchaeales Nanobdellota* tips on the other side. The same is true of every other named multi-tip *Nanobdellota* order in our sampling: *Pacearchaeales*, *Maxwellarchaeales*, *Nanobdellales*, *Jingweiarchaeales*, *Tiddalikarchaeales*, CSSed11-243R1, UBA10117, and WJKC01 in both trees, with *Parvarchaeales* (58 tips) and JAPDLS01 (10 tips) additionally resolved as single bipartitions at the 1,239-tip ar71 scale where they were not at the smaller 444-tip ar53 scale. The two inference substrates differ in scope and resolution. ar53 is a broadly-trained archaea-wide marker set used by GTDB for phylum-level placement across all Archaea; ar71 is a *Nanobdellota*-specific 71-marker set built from our 208 complete genomes (Results *marker set*; Methods *ar71 phylogeny*) and inferred over 1,239 taxa to maximise within-*Nanobdellota* resolution. They share only a subset of canonical informational markers. Their agreement on order-level monophyly across this scope–resolution gradient is the strength of the result.

The within-phylum resolution available at this sampling extends Liu et al. (2018), who identified 26 *Woesearchaeota* subgroups (Woese-1 through Woese-26) but tested monophyly within the order rather than across the order-level *Nanobdellota* taxonomy. Our complete-genome dataset, embedded in the 1,239-taxon ar71 tree with R232 MAG context, supports the existing GTDB R232 order-level circumscription of *Nanobdellota* — *Pacearchaeales* and *Maxwellarchaeales* (= GTDB R232 *SCGC-AAA011-G17*) each form a single bipartition in both primary inferences and retain their order rank. We therefore propose no change to the rank of any *Nanobdellota* order in this paper, but propose a four-rank SeqCode nomenclatural chain that retires the GTDB *SCGC-AAA011-G17* placeholder name (*Taxonomic proposal*: *Maxwellarchaeales* ord. nov., *Maxwellarchaeaceae* fam. nov., *Maxwellarchaeum* gen. nov., *Maxwellarchaeum balticum* sp. nov.); the relationship of the *Nanobdellota* orders to the broader DPANN superphylum, including the Baker et al. (2025) inter-phylum reconstruction, is at a different scale and is unaffected by the within-phylum monophyly result reported here.

### Synthesis

Two findings emerge from the 208-complete-genome substrate: a within-phylum metabolic gradient across the three environmental *Nanobdellota* orders, and recurrent archaea-to-CPR transfer of Form III RuBisCO at the same orders that drive the metabolic gradient. *First*, the within-phylum metabolic gradient (*Woesearchaeales* lineages retaining partial glycolysis and a V/A-type ATPase; *Pacearchaeales* and *Maxwellarchaeales* lineages retaining only Form III RuBisCO, PEP synthase, and ferredoxin on top of an otherwise universal information-processing core) is a gradient *within* a small-genome phylum, not a contrast between it and a larger-genome relative; it sits within rather than across the DPANN size spectrum and constitutes an evolutionary observation that any larger-scale account of DPANN evolution, including the reductive-from-free-living scenario advanced by Baker et al. (2025), must accommodate regardless of which inter-phylum ancestor scenario eventually proves correct. *Second*, Form III RuBisCO — the most distinctive metabolic feature of the small-genome *Pacearchaeales* and *Maxwellarchaeales* lineages — is also the locus of recurrent inter-kingdom transfer: a 4,262-tip rbcL gene tree identifies nine candidate archaea-to-*Patescibacteriota* transfer events across our sampled environments, including one to a co-sampled Baltic *Minisyncoccia* (the CPR-equivalent class containing the recently cultured archaea-parasitizing isolate *Minisyncoccus archaeiphilus*; Nakajima et al. 2025), against only two candidate reverse-direction transfers (both below UFBoot 60). The qualitative asymmetry between the candidate counts is reported as a directional pattern consistent with archaea-to-CPR being the more frequently identified direction in our data rather than as a calibrated rate. rbcL transfer therefore further blurs the metabolic boundary between archaeal and bacterial small-genome lineages at exactly the orders where the within-phylum metabolic divide is sharpest, locating a load-bearing carbon-metabolism enzyme in the inter-kingdom contact zone the metabolic-gradient finding describes.

### Taxonomic proposal

The SeqCode (Hedlund et al. 2022) is a parallel nomenclature framework to the International Code of Nomenclature of Prokaryotes that permits the formal naming of prokaryotic taxa from genome sequences alone, without a cultivated type strain. Each protologue below comprises a Latin etymology, a diagnosis (the phenotypic and genotypic features that circumscribe the taxon), a designated type genome (the nomenclatural reference per SeqCode rules; an uncultivated type strain is not required), and a set of distinguishing features that separate the taxon from its closest relatives.

We propose four taxa under the SeqCode at order, family, genus, and species rank, anchored on a complete-genome type from this study, replacing the GTDB-recognized placeholder order *SCGC-AAA011-G17* with a Latin-binomial nomenclatural chain: *Maxwellarchaeales* ord. nov., *Maxwellarchaeaceae* fam. nov., *Maxwellarchaeum* gen. nov., and *Maxwellarchaeum balticum* sp. nov. The order-level circumscription is unchanged: the *Maxwellarchaeales* clade (= GTDB R232 *SCGC-AAA011-G17*) is recovered as a single monophyletic bipartition in our ar53 and ar71 phylogenies (Discussion *Woesearchaeales is recovered monophyletic*); the present proposal does not alter its rank, it replaces the GTDB placeholder name with a Latin name.

### Maxwellarchaeales ord. nov

*Etymology* — Max.well.ar.chae.a’les. N.L. neut. n. *Maxwellarchaeum*, the type genus of the order;-ales, the ending to denote an order; N.L. fem. pl. n. *Maxwellarchaeales*, the order of *Maxwellarchaeum*. The order honors James Clerk Maxwell (1831–1879) through its type genus.

*Diagnosis* — An order of ultrasmall, uncultivated archaea within the phylum *Nanobdellota* (superphylum DPANN), recovered as a single monophyletic bipartition in both the ar53 supermatrix (444-tip ML tree from a 446-taxon alignment) and the *Nanobdellota*-tuned ar71 supermatrix (1,239 taxa) presented in this study. Members are environmentally sampled aquatic archaea characterized by reduced genomes (719 kbp to 1,473 kbp across the 31 complete genomes from this study; median 1,030 kbp), GC content 26–42% (median 31%), absence of a functional electron transport chain and V/A-type ATPase, and a metabolic capability reduced to an information-processing core plus Form III RuBisCO (KEGG K01601), PEP synthase, and ferredoxin where annotated. Form III RuBisCO is carried by 87% of complete genomes (27/31). Replication initiation is via archaeal ORC1/Cdc6 (KEGG K10725); no bacterial dnaA is detected. The order corresponds to the GTDB R232 placeholder order *SCGC-AAA011-G17*.

*Maxwellarchaeales* is distinguished from its sister metabolically-reduced order *Pacearchaeales* (54 complete genomes from this study) by larger median genome size (1,030 vs 739 kbp), lower characteristic GC content (median 31% vs 34%), retention of pyruvate:ferredoxin oxidoreductase in a subset of members (α 17/31, β 13/31, δ 13/31, γ 11/31 vs 3/54, 3/54, 3/54, 0/54 in *Pacearchaeales*), and succinyl-CoA synthetase α-subunit in 6/31 versus 0/54 in *Pacearchaeales*.

*Type family* — *Maxwellarchaeaceae* fam. nov. (described below).

### Maxwellarchaeaceae fam. nov

*Etymology* — Max.well.ar.chae.a.ce’ae. N.L. neut. n. *Maxwellarchaeum*, the type genus of the family;-aceae, the ending to denote a family; N.L. fem. pl. n. *Maxwellarchaeaceae*, the family of *Maxwellarchaeum*.

*Diagnosis* — Currently the sole named family within order *Maxwellarchaeales*; circumscription identical to that of the order. The family contains all 31 complete *Maxwellarchaeales* genomes presented in this study and is open to subsequent subdivision as additional named genera are described from the lineage.

*Type genus* — *Maxwellarchaeum* gen. nov. (described below).

### Maxwellarchaeum gen. nov

*Etymology* — Max.well.ar’chae.um. N.L. neut. n. *Maxwellarchaeum*, honoring James Clerk Maxwell (1831–1879); in conjunction with N.L. neut. n. *archaeum* (from Gr. masc. n. *archaios*, ancient), an archaeon; the compound is treated as a neuter noun. *Maxwellarchaeum* is here proposed at genus rank for the first time under the SeqCode.

*Diagnosis* — A genus of ultrasmall, uncultivated archaea within family *Maxwellarchaeaceae* fam. nov. (described above), order *Maxwellarchaeales* ord. nov. (described above), phylum *Nanobdellota*, superphylum DPANN. Genome size 719 kbp to 1,473 kbp across the 31 complete *Maxwellarchaeales* genomes from this study (median 1,030 kbp); GC content 26–42% (median 31%); Form III RuBisCO (KEGG K01601) carried by 87% of complete genomes (27/31), a frequent but not strictly diagnostic feature; replication initiation via archaeal ORC1/ Cdc6 (KEGG K10725) with no detectable bacterial dnaA; no functional electron transport chain or V/A-type ATPase encoded; metabolic capability reduced to an information-processing core plus Form III RuBisCO, PEP synthase, and ferredoxin where annotated.

*Maxwellarchaeum* is distinguished from the 54 complete *Pacearchaeales* genomes from this study (the sister metabolically-reduced order in our complete-genome dataset) by larger median genome size (1,030 vs 739 kbp), lower characteristic GC content (median 31% vs 34%), retention of pyruvate:ferredoxin oxidoreductase in a subset of members (α 17/31, β 13/31, δ 13/31, γ 11/31 *Maxwellarchaeales* genomes versus 3/54, 3/54, 3/54, 0/54 *Pacearchaeales* genomes), and succinyl-CoA synthetase α-subunit in 6/31 *Maxwellarchaeales* genomes versus 0/54 *Pacearchaeales* genomes.

*Type species* — *Maxwellarchaeum balticum* sp. nov. (described below).

### Maxwellarchaeum balticum sp. nov

*Etymology* — bal’ti.cum. L. neut. adj. *balticum*, pertaining to the Baltic Sea; the type genome was sequenced from a Baltic Sea water-column metagenome.

*Diagnosis* — Type species of *Maxwellarchaeum* gen. nov. A complete archaeal genome (ORC1-rotated) assembled from a Baltic Sea water-column metagenome (sample B10). Genome size 842,781 bp; GC content 31%; coding density ∼93%; 999 predicted protein-coding genes (Pyrodigal v3.6.3, single mode). CheckM2 (v1.1.0) completeness 95.47%, contamination 0.14%. Carries Form III RuBisCO (rbcL, KEGG K01601) at strict KofamScan threshold and the archaeal replication initiation protein ORC1/Cdc6 (KEGG K10725); no detectable bacterial dnaA, no functional electron transport chain, no V/A-type ATPase, no F-type ATP synthase. Recovered as a member of the *Maxwellarchaeales* clade in the ar71 phylogeny on 1,239 taxa. Selected as the type genome by the lowest composite score (lower = better; Methods *Type-genome composite ranking*) among the 17 candidate *Maxwellarchaeales* genomes (the *qualified subset*) that passed our ≥20× coverage filter; the composite is the sum of five min-max-normalized components, each oriented so lower = better: CheckM2 completeness, contamination, GC distance from the median of the 17-genome qualified subset, relative size distance from the same subset’s median, and mean patristic distance to other *Maxwellarchaeales* tips in the ar71 ML tree.

B10_u40295554 ranks 1st of 17 with composite 0.603 (driven by exact match to the qualified-subset median genome size of 842.8 kbp, GC of 31% matching the qualified-subset median, completeness 95.47%, contamination 0.14%, and Depth1 sequencing coverage of 194×); the next-ranked candidate (B7_u10090349) is at composite 0.625, comparable enough that it could substitute as type genome under SeqCode rules without altering the genus circumscription. The 842.8 kbp qualified-subset median (17 genomes) is used for the type-genome composite-ranking score; the 1,030 kbp order-level median reported in the *Maxwellarchaeum* genus diagnosis is computed across all 31 complete *Maxwellarchaeales* genomes from this study.

*Pairwise ANI to closest available reference* — The Maxwell type genome was screened against all 28 other complete *Maxwellarchaeales* genomes from this study (skani v0.3.1 dist) and against the GTDB R232 high-quality *SCGC-AAA011-G17* reference set (the GTDB placeholder for the same clade). Within our complete-genome set, B5_u18733786 (99.57% ANI; reference alignment fraction 93.14%, query 92.15%) and B7_u10090349 (99.23% ANI) fall above the 95% same-species cut-off and are members of the same species as the type. The closest non-conspecific match within our 28 is SA1420A_u24606309 at 83.82% ANI (alignment fraction 20.2%), well below the 95% species-distinguishability threshold under SeqCode Rec. 2 and below skani’s default screen-coverage requirement; the remaining 25 complete *Maxwellarchaeales* genomes fall below the skani default screen entirely (∼80% ANI). The closest non-conspecific neighbour from the GTDB R232 reference pool is GCA_018901245.1 at 85.13% ANI. The species *Maxwellarchaeum balticum* therefore comprises at minimum the type genome plus B5_u18733786 and B7_u10090349, all from Baltic Sea water-column samples in the present study.

*Type genome* — B10_u40295554, a complete genome from sample B10 (Baltic Sea water column), Oxford Nanopore PromethION sequencing, myloasm v0.4.0 assembly, rotated to ORC1/Cdc6 at position 1 with UniRef90 verification. Sequencing coverage at ≥99% identity: 194× (myloasm Depth1). As the lineage is uncultivated, the type genome serves as the nomenclatural type per SeqCode rules (Hedlund et al. 2022); raw reads for the source B10 metagenome are deposited under NCBI BioProject `PRJNA1461195` (BioSample `SAMN57849840`); the type genome is deposited at GenBank as `JBYEVE000000000` (per-genome BioSample `SAMN59124702`).

### Outstanding nomenclatural items

The names *Maxwellarchaeales*, *Maxwellarchaeaceae*, *Maxwellarchaeum*, and *Maxwellarchaeum balticum* will be registered with the SeqCode Registry at https://seqco.de/. Where possible, the registration identifiers will be added to the published version; otherwise, the publication DOI will be entered into the Registry post-publication to complete validation under the SeqCode (Hedlund et al. 2022).

## Methods

Filenames in monospace throughout this paper (e.g., `custom_hmms/ar71/`) refer to subdirectories of the Zenodo archive accompanying this paper (DOI: 10.5281/zenodo.20174424; enumerated in *Using the resource*). Each script that produced a paper figure or table is included in that archive under `scripts/`.

### Sample collection and sequencing

Metagenomes were obtained from six samples spanning two environments. Four Baltic Sea water-column samples (B5, B7, B10, B14) were collected by Stefan Bertilsson on a Baltic cruise. Two Fennoscandian groundwater samples from the Äspö Hard Rock Laboratory — KR0015B (69 m below sea level) and SA1420A (201 m below sea level) — were collected by Mark Dopson. DNA extraction and Oxford Nanopore library preparation followed the protocol described in Lui and Nielsen (2024). Sequencing libraries were prepared with the Oxford Nanopore SQK-LSK114 ligation sequencing kit and run on PromethION R10.4.1 flow cells. Basecalling was performed with dorado v0.9.6.

### Assembly and quality filtering

Metagenome assembly was performed with myloasm v0.4.0 (Shaw et al. 2026) with default parameters. Assembled contigs ≥ 500 kbp were assessed with CheckM2 v1.1.0 (Chklovski et al. 2023) with the `uniref100.KO.1.dmnd` reference database (downloaded from the CheckM2 distribution on 2024-12-15; the same.dmnd build was used across all genus papers in this series for cross-paper consistency).

High-quality genomes were defined as single-contig assemblies with ≥ 90% completeness and < 5% contamination. Circular (complete) genomes were identified from the assembly metadata.

### Taxonomic classification

All high-quality contigs were classified with GTDB-Tk v2.6.1 (Chaumeil et al. 2022) against GTDB release R226, using the archaeal ar53 marker set. *Nanobdellota* genomes were identified by filtering for phylum p Nanobdellota. *Patescibacteriota* genomes co-assembled from the same six samples (B5, B7, B10, B14, KR0015B, SA1420A) and passing the same quality filters (single-contig, ≥90% CheckM2 completeness, <5% contamination, GTDB-Tk classification to phylum p Patescibacteriota) numbered 632; these were used as the bacterial co-occurrence set for the rbcL transfer analysis (*Form III rbcL transfer from archaea to CPR*).

### Rotation to ORC1/Cdc6

Circular genomes were rotated to place ORC1/Cdc6 at position 1 using an in-house rotation script. For each genome, Pyrodigal v3.6.3 (Larralde 2022; single mode) predicted genes on the unrotated sequence, and all predicted proteins were searched against a curated ORC1 reference database of 19 sequences via MMseqs2 v18.8cc5c (Steinegger and Söding 2017) easy-search (sensitivity 7.5, e-value threshold 10⁻⁵, max-seqs 1). The ORC1 reference was constructed from Baltic genomes by searching predicted proteins against UniRef90 (release 2024_02) and extracting those annotated as “ORC1-type DNA replication protein” or “cell division control protein 6.” The best full-length hit (query/target length ratio ≥ 0.80) was selected as the rotation target. Rotation placed the ORC1 start codon at position 1 on the forward strand, with reverse-complementation applied when ORC1 was on the minus strand. Each rotated genome was verified by re-running gene prediction and confirming that gene_1 matched the ORC1 reference. Per-genome ORC1 verification (gene_1 identity, start position, strand, query coverage, e-value vs the ORC1 reference, UniRef90 best-hit description) is recorded in `orc1_rotation/ orc1_full_audit.tsv`. Rotation is a processing step rather than a completeness criterion: a complete genome (single-contig, assembler-circular) is included in the cohort regardless of whether its ORC1/Cdc6 can be located by the rotation pipeline, since failure to identify a hit may reflect a frameshifted ORC1 (cf. the 1,214 frameshift events documented across the cohort, *Frameshift detection*) rather than genuine absence.

### Gene prediction and orthogroup inference

Genes for the rotated complete genomes were predicted at the gene-prediction step of the in-house rotation pipeline (Pyrodigal v3.6.3, single mode, `force_nonsd=False`); the canonical proteome is the union of `genes/<genome>.faa`. We use `force_nonsd=False` rather than the project-default `force_nonsd=True` because rotation places the ORC1/ Cdc6 start codon at position 1 with no upstream context, and empirical testing showed that under `force_nonsd=True` Pyrodigal’s non-SD start-codon scoring privileges upstream ATGs in close proximity to the highest-scoring frame, which in the absence of upstream context (positions <0 lie wrapped past the chromosome end) selects an ATG 6–9 nt upstream of the conserved ORC1 N-terminus and extends the gene_1 prediction beyond the reference (verified against the reference ORC1 set). The setting is specific to this rotated-input pipeline and does not reflect a general SD-vs-non-SD choice for archaea.

Single-copy orthogroups across the complete-genome set were inferred with OrthoFinder v3.1.4 (Emms and Kelly 2019) using all-vs-all DIAMOND (Buchfink et al. 2021; OrthoFinder-bundled binary) search (default parameters), MCL clustering with inflation 1.2 (OrthoFinder default), MAFFT for orthogroup alignment, and IQ-TREE3 for gene tree inference: `orthofinder-f`

`<genomes>-t 64-a 32-M msa-A mafft-T iqtree3`. Single-copy orthogroups were selected by OrthoFinder’s automatic `DetermineOrthogroupsForSpeciesTree` algorithm (an internal OrthoFinder helper, defined in the OrthoFinder distribution at `scripts_of/trees_msa.py`; algorithm cited as Emms and Kelly 2019) which walks a single-copy fraction threshold from 1.0 downward — applying a probabilistic test for hidden paralogy at each step — and stops when at least 100 orthogroups qualify and further relaxation no longer increases the count by ≥2× per relative decrease in the threshold (`nOGsMin = 100`, `increase_required = 2.0`).

For our 206-genome run the algorithm halted at a single-copy-fraction threshold of approximately 96.1% (the smallest threshold at which the 100-OG count and ≥2× growth criterion were both satisfied) and emitted 101 selected orthogroups; OrthoFinder writes the 101-OG list to `Results_Apr15/Species_Tree/ Orthogroups_for_concatenated_alignment.txt`. The 96.1% figure is the OF stopping threshold, not the realized per-OG occupancy: per-OG single-copy rates across the 101 selected OGs span 19.4–100% (43 of the 101 OGs sit below the 96.1% threshold and are retained because removing them would drop the OG count below the algorithm’s `nOGsMin = 100` floor). We use this canonical OF-selected set throughout. The list with per-OG occupancy statistics is reproduced by `phylogeny/ scripts/derive_sc_ogs.py` to `orthofinder/ single_copy_orthogroups_206.tsv`.

The 206-genome complete-cohort snapshot used for OrthoFinder seed inference was the cohort current at the run timestamp; the cohort grew to 208 when two genomes whose ORC1 rotation could not be confirmed at that snapshot were re-evaluated and re-included under the cohort definition (*Rotation to ORC1/Cdc6*). The 71 calibrated markers are unaffected — calibration was against the 1,288-genome R232 pool, not the 206-genome OF input — and the 2 additional complete genomes were screened against the calibrated GA thresholds before joining the 446-taxon downstream set. We retain the 206-genome OF run-of-record (`Results_Apr15`) rather than re-running OrthoFinder on 208, since the 101 selected orthogroups are stable to a 1% cohort perturbation and the marker set is the load-bearing output for the phylogeny.

### Functional annotation and crosscheck

KofamScan v1.3.0 was run on all 274,837 proteins from the 208 complete genomes and separately on 266,741 NCBI proteins, using KEGG HMM profiles downloaded 27 February 2026 paired with the corresponding `ko_list` score-threshold file downloaded 24 March 2026 (KEGG profile-specific bit thresholds are static once published, so the 25-day download gap reflects asynchronous local re-downloads of the same KEGG release rather than a release-version mismatch; the threshold values applied here are those carried in the 24 March `ko_list`). Strict profile-specific score thresholds were applied for all metabolic claims. The crosscheck (`scripts/ kofamscan_crosscheck_v2.py`) compared KofamScan strict and relaxed annotations against UniRef90 best-hit descriptions from MMseqs2 v18.8cc5c easy-search (sensitivity 7.5, e-value threshold 10⁻³ — more permissive than the 10⁻⁵ used for ORC1 identification, to capture divergent homologs, max-seqs 1, format: query, target, fident, evalue, bits, qlen, tlen, alnlen, raw, theader). A KO was classified as “present” if detected at strict threshold with UniRef90 confirmation, “present (divergent)” if at relaxed threshold (0.5–1.0× profile score) with UniRef90 confirmation, “partial” if in a subset of genomes, or “absent” if below 0.5× threshold or without UniRef90 confirmation. KO-to-gene-name mappings were loaded from the canonical ko_list file at runtime; no identifier labels were hand-typed.

### ANI-based species clustering

Pairwise average nucleotide identity (ANI) was computed with skani v0.3.1 (Shaw and Yu 2023) using the dist subcommand with default parameters (screen threshold s=80, no minimum alignment fraction filter) for all pairwise comparisons. Species-level clusters were defined at 95% ANI by single-linkage clustering of connected components (`scripts/cluster_species_95ani.py`, Union-Find on the skani output).

### NCBI comparison

All *Nanobdellota* assemblies (870 after GCA/GCF deduplication) were downloaded from NCBI on 12 April 2026 using the datasets CLI v18.10.2. CheckM2 v1.1.0 was run on all 870 genomes without a size threshold. GTDB taxonomy for the 245 high-quality genomes was obtained by two methods: 111 genomes present in the GTDB R226 reference were looked up directly from the reference taxonomy file (gtdb_taxonomy.tsv); the remaining 134 were classified with GTDB-Tk v2.6.1 classify_wf. Gene prediction was performed with Pyrodigal v3.6.3 (force_nonsd=True) on the 238 confirmed *Nanobdellota* genomes, yielding 266,741 proteins. The `force_nonsd` flag differs between our 208 (False) and the NCBI 238 (True) because our complete-genome cohort is rotated to ORC1/Cdc6 at position 1 with no upstream context — under `force_nonsd=True` Pyrodigal’s start-codon scoring then over-extends the gene_1 prediction by 6–9 nt; the NCBI MAGs are not rotated and retain natural upstream context, so the project-default `force_nonsd=True` is used there. The residual between-set difference is an N-terminal length difference of ≤9 nt for a small subset of cross-set homologous genes (predominantly those with ORC1-like 5′ UTR motifs); the per-gene KO recovery underlying our metabolic claims and the genome-size–KO correlations is not affected by this end-effect because KO calls are HMM matches against full-length conserved domains rather than start-codon positions.

### Marker set selection and HMM construction

The *Nanobdellota* marker set ar71 used here was constructed from the 101 OrthoFinder single-copy orthogroups by an iterative HMM-refinement procedure with calibrated per-marker score thresholds.

The construction is a general-purpose *Nanobdellota* phylogenomic instrument rather than a paper-specific artefact: the same 71-marker set is used in this paper for the within-*Nanobdellota* phylogeny and is reusable as-is for any future *Nanobdellota* phylogenomic analysis. The deposited HMMs (with embedded GA / NC bit-score thresholds) are the only inputs needed; no per-paper tuning is involved. For each candidate orthogroup, an initial profile HMM was built with HMMER v3.4 (Eddy 2011) `hmmbuild` from the 206-genome single-copy alignment, then expanded by aligning high-scoring hits from the 1,288-genome *Nanobdellota* calibration pool (the 257 our-Nano stems in the OF-era proteome together with 1,031 GTDB R232 high-quality *Nanobdellota* MAGs) into the alignment with MAFFT v7.525 (Katoh and Standley 2013) `--add --keeplength` and rebuilt with `hmmbuild`. Iteration continued until the single-copy hit set converged on the 1,288-genome pool or eight refinement rounds were reached, and each marker was rolled back to the iteration that maximized single-copy recovery (some markers degraded with continued iteration as paralogs accumulated; rollback was applied per-marker). Hits were accepted only at envelope coverage ≥70% of HMM length to drop partial / fragment matches that inflate apparent paralogy. Per-marker bit-score thresholds (HMMER GA / NC) were calibrated against 206-genome OrthoFinder OG-membership ground truth (TPs = 206-genome OG members; FPs = all other 206-genome proteins): GA was set to the midpoint of min(TP) and max(FP) when the two distributions were separable, otherwise to the 1st percentile of TPs; NC was set to max(FP). The calibrated HMM was then searched against the 1,288-genome pool with `hmmsearch --cut_ga` and three quality criteria evaluated: single-copy recovery (target ≥1,100/1,288), multi-copy rate (target ≤5%), and GA value (target ≥30 bits). 71 of the 101 candidate orthogroups satisfied all three criteria; the remaining 30 were excluded for paralog distribution across *Nanobdellota* subsets, under-detection in distantly-related orders, or lack of a clean GA boundary between true positives and false positives. The 71-marker ar71 set is deposited at `custom_hmms/ar71/` together with per-marker metadata (`custom_hmms/ar71/index.tsv`) and the calibrated HMMs (`custom_hmms/ar71/hmms/<nnn>_`

`<marker>.hmm`). The 71-marker set was compared against the GTDB archaeal marker sets ar53 (GTDB R226) and ar122 (Parks et al. 2018) to quantify overlap; OrthoFinder OG identifiers for the 71 ar71 markers are recorded in `custom_hmms/ ar71/index.tsv`.

### Marker-set characterization alignments

For each of the 71 ar71 markers, single-copy amino acid sequences were extracted from all 446 taxa (208 complete genomes — 206 by OrthoFinder orthogroup membership and 2 by hmmsearch under the calibrated GA threshold; 238 NCBI MAGs by hmmsearch under the same calibrated GA threshold) and aligned with MAFFT v7.525 L-INS-i (`--localpair --maxiterate 1000 --thread 4`). Each alignment was trimmed with BMGE v2.0 (Criscuolo and Gribaldo 2010) using the BLOSUM30 substitution matrix and default entropy/gap settings (`-t AA-m BLOSUM30`). BLOSUM30 was selected over the BMGE default BLOSUM62 following the Criscuolo & Gribaldo (2010) recommendation of BLOSUM30 for distantly-related sequences: within-phylum median pairwise identity ranged from 31.4% to 65.9% across the marker set, with some pairs as low as 13.2%, and at these divergence levels BLOSUM62 over-trims phylogenetically informative positions. Within-phylum identity statistics reported in Discussion *marker set* are computed across these 71 trimmed alignments. The same alignment and trimming procedure (with BLOSUM30) was used at the larger scale for the ar71 phylogeny on 1,239 taxa (*ar71 phylogeny*).

### Phylogenomic inference

The ar53 marker alignment was extracted for each of the 446 taxa either from GTDB-Tk output (for genomes classified by GTDB-Tk) or from the GTDB R226 reference MSA (for genomes already in the reference), with the GTDB R226 column mask applied to yield an alignment of 8,062 amino acid positions; IQ-TREE collapsed two identical sequences before inference, leaving 444 unique tips in the resulting ML tree. The ar53 phylogeny was inferred with IQ-TREE v3.0.1 (Wong et al. 2025) from the unpartitioned supermatrix under ModelFinder Plus with model selection constrained to substitution matrices LG, Q.pfam, WAG, JTT, VT, rtREV and rate-heterogeneity models G, I+G, R2, R4, R6, R8, R10 (`--mset LG,Q.pfam,WAG,JTT,VT,rtREV--mrate G,I+G,R2,R4,R6,R8,R10 --cmax 10`): `iqtree3-s ar53_444.fasta-m MFP-B 1000-alrt 1000-T 32`. The ar53 run was kept unpartitioned to match standard GTDB methodology. The run used 1,000 ultrafast bootstrap replicates (UFBoot; Hoang et al. 2018) and 1,000 SH-aLRT replicates (Guindon et al. 2010). The consensus tree was midpoint-rooted with ete4 v4.4.0 (Huerta-Cepas et al. 2016; ete4 is the v4 successor to ete3 cited there) (`Tree.get_midpoint_outgroup` + `set_outgroup`). The chi-squared composition test failed for most sequences — an expected consequence of DPANN AT-biased compositional heterogeneity rather than a data-quality issue. The *Nanobdellota*-specific phylogeny is described separately in *ar71 phylogeny*. The radial phylogeny was rendered with ggtree v4.1.2 (Yu 2022) in R. The unconstrained ar53 ML topology is at `phylogeny/tree.contree`.

### ar71 phylogeny

The ar71 marker set (Methods *Marker set selection and HMM construction*) was searched (HMMER v3.4 `hmmsearch --cut_ga`) against the predicted proteomes of 1,031 GTDB R232 high-quality *Nanobdellota* genomes plus our complete-genome set, with the calibrated per-marker GA bit-score thresholds. Single-copy hits passing GA were extracted, aligned per marker with MAFFT v7.525 L-INS-i (`--localpair--maxiterate 1000`), trimmed with BMGE v2.0 (Criscuolo and Gribaldo 2010) under BLOSUM30, and concatenated end-to-end into a single-block supermatrix of 1,239 taxa × 11,883 amino acid columns. Per-marker BMGE-trimmed alignment lengths and per-tip occupancy are recorded in `ar71_tree/marker_lengths.tsv` and `ar71_tree/ tip_occupancy.tsv`; the partition definition `ar71_tree/partitions.txt` is reference metadata only — inference is unpartitioned. Phylogenomic inference used IQ-TREE v3.1.1 with the LG+F+R10 model (`-m LG+F+R10`), 1,000 ultrafast bootstrap replicates, 1,000 SH-aLRT replicates, and explicit thread count `-T 64`. The choice of a single LG+F+R10 model rather than a per-marker partitioned scheme was deliberate: at the within-*Nanobdellota* scale (1,239 deeply-divergent tips), partitioned schemes inflate per-partition rate-heterogeneity parameter variance and produced lower bootstrap convergence than the single-model alternative in our pilot inference. UFBoot split correlation reached 0.988 at the 1,000-iteration cap, below IQ-TREE’s default `-bcor 0.99` cutoff (a workflow convention, not a quality threshold) but consistent with stable per-branch bootstrap support across the final ∼200 iterations. The tree consumed by paper-internal analyses is `ar71_tree/ ar71.treefile`. The order-level monophyly conclusions used downstream are independently recovered by the parallel ar53 supermatrix (which reached UFBoot ≥ 0.99) on a different marker set at smaller taxon scope.

### Per-order monophyly assessment

For each named *Nanobdellota* order with two or more taxa, monophyly was assessed in both the ar53 phylogeny (444-tip ML tree from a 446-taxon alignment) and the ar71 phylogeny (1,239 taxa) by checking whether the order’s tips form one side of a single bipartition in the inferred unrooted tree (`phylogeny/scripts/ check_unrooted_monophyly.py`). The unrooted bipartition test is rooting-independent: a set S of tips is monophyletic in an unrooted tree T iff there exists an edge of T whose removal partitions the tips into exactly {S, V}. For each order, every node n in the tree was examined and its `tips_below(n)` compared against the target set; the order was deemed monophyletic if any node n satisfied `tips_below(n) == S` or `tips_below(n) == V\S`. In the 444-tip ar53 tree, every named multi-tip *Nanobdellota* order — *Woesearchaeales* (205 tips), *Pacearchaeales* (83), *SCGC-AAA011-G17* (73; here renamed *Maxwellarchaeales*; *Taxonomic proposal*), *Nanobdellales* (16), *Jingweiarchaeales* (12), CSSed11-243R1 (9), *Tiddalikarchaeales* (7), UBA10117 (7), WJKC01 (6) — is monophyletic by unrooted bipartition; *Parvarchaeales* (11 tips) and JAPDLS01 (11 tips) are not. In the 1,239-taxon ar71 tree, all 13 multi-tip *Nanobdellota* orders, including *Woesearchaeales* (728 tips), *Parvarchaeales* (58 tips), and JAPDLS01 (10 tips), are monophyletic by unrooted bipartition.

### Type-genome composite ranking for SeqCode protologues

The type genome for the *Maxwellarchaeum balticum* SeqCode protologue was selected from a candidate set restricted to *Maxwellarchaeales* genomes that passed a ≥20× sequencing-coverage filter on Depth1 (myloasm read-mapping at ≥99% identity): 17 candidates after the filter (`phylogeny/scripts/ maxwellarchaeae_type_genome_candidates.tsv`). For each candidate five components were computed: (1) CheckM2 completeness; (2) CheckM2 contamination; (3) GC-content distance from the qualified-subset median |GC − median|; (4) relative genome-size distance from the qualified-subset median |size − median| / median; and (5) phylogenetic centrality, the mean patristic distance from this tip to all other qualified-subset tips in the ar71 ML tree (`ar71_tree/ ar71.treefile`). Each component was min-max-normalized to [0, 1] within the qualified candidate set, oriented so that lower = better on every dimension (completeness mapped 100 to 0, all other components have natural lower-is-better orientation). The composite score is the *sum* of the five normalized components — range 0–5, lower = better, with 0 representing the (in general unattainable) joint optimum on all five dimensions. The full per-candidate score table is written to the candidate TSV (columns `n_completeness`, `n_contamination`, `n_gc_dist`, `n_size_dist`, `n_phylo_dist`, `score`, `rank`). Reproduction script: `phylogeny/ scripts/rank_type_genome_maxwell.py`. B10_u40295554 was rank 1 of 17 *Maxwellarchaeales* candidates with composite 0.603 (next-ranked B7_u10090349 at 0.625; Δ 0.022). The narrow Δ at the top of the table indicates that the *Maxwellarchaeum balticum* type-genome choice is robust at rank 1 but the rank-2 candidate is comparable and could substitute under SeqCode rules without altering the genus circumscription.

### rbcL phylogeny

The rbcL gene tree was constructed from seven sources of strict-threshold K01601 hits, with hit counts (search-space genomes shown in parentheses): (i) `our_nano`, 247 hits across the union of our 256 internally-assembled *Nanobdellota* proteomes (133 hits, comprising 105 hits from 95 distinct genomes among the 208 complete; the 10-hit margin is rbcL paralogs in the same genome; plus 28 hits from the 48 high-quality non-circular) and the 238 high-quality NCBI *Nanobdellota* MAGs (114 hits), for 494 search-space genomes in total; (ii) `our_cpr`, 42 hits in our 632 co-sampled high-quality *Patescibacteriota* proteomes from the same six samples; (iii) `r232_nano`, 341 hits in 1,031 GTDB R232 high-quality *Nanobdellota* proteomes; (iv) `r232_arch`, 3,680 hits in 8,776 R232 high-quality non-*Nanobdellota* archaeal proteomes; (v) `r232_cpr`, 372 hits in 11,567 R232 high-quality *Patescibacteriota* proteomes (after Pyrodigal v3.6.3 gene prediction on the downloaded genomes); (vi) `r232_form3_bact`, 17 hits in 317 R232 high-quality Form III bacterial genomes (Hydrothermota, Thermodesulfobiota, and other phyla flagged for Form III rbcL); (vii) `KEGG`, 1,498 K01601 reference entries via the KEGG `/link/genes/ko:K01601` API (downloaded 27 February 2026, the same release date as the KofamScan profiles). After cross-source dedup (1,921 duplicates dropped) the combined set was 4,276 unique sequences; the IQ-TREE-deduplicated alignment yielded 4,262 tips in the inferred tree. Sequences were aligned with MAFFT v7.525 L-INS-i (3,036 columns at the pre-trim stage, reflecting the broad insertion heterogeneity of an alignment spanning Form IB plants, Form III archaea, Form III bacteria, and Form-IV-like RuBisCOs), trimmed with BMGE v2.0 under BLOSUM30 (296 amino-acid positions retained, 9.7% of pre-trim, since the cross-form heterogeneity drops most pre-trim columns; the 296 retained columns are those with informative coverage across the full sequence pool), and inferred with IQ-TREE v3.0.1 under model selection constrained to LG with FreeRate rate heterogeneity at 6, 8, or 10 categories (`--mset LG --mrate R6,R8,R10 -- cmax 10`), 1,000 ultrafast bootstrap replicates, 1,000 SH-aLRT replicates (`-T 32`). The strict K01601 score threshold used to source all archaeal and bacterial Form III sequences was the profile-specific bit threshold from the KofamScan ko_list (310.97 bits for K01601). The `r232_form3_bact` pool was defined as GTDB R232 high-quality (≥90% completeness, <5% contamination) genomes from any phylum *outside* p Patescibacteriota and outside p Nanobdellota that contained at least one strict-threshold K01601 hit; this captures the historically-described Form III–bearing bacterial phyla (notably Hydrothermota and Thermodesulfobiota) as well as any further Form III–bearing bacterial genomes regardless of phylum assignment, and totalled 317 genomes contributing 17 hits. The constrained model selection rationale: a prior exhaustive ModelFinder run on an earlier, smaller rbcL alignment selected LG+R10 as BIC-best; restricting the present run to the same family of models reflects this prior evidence while still testing across rate-heterogeneity dimensionality. The final tree’s UFBoot split-correlation reached 0.922, below IQ-TREE’s `-bcor 0.99` workflow default (a workflow convention, not a quality threshold) but consistent with stable per-branch bootstrap support; the value is treated as a soft caveat in interpretation, see Discussion *RuBisCO*.

The tree was post-hoc pruned for tractable visualization and cross-kingdom analysis (`rubisco_tree_v2/prune_rbcl_v2.py`): Strategy A dereplicated paralogs by retaining the highest-scoring rbcL per genome / KEGG species (4,262 to 3,888 tips); Strategy B walked the tree to identify maximal subtrees containing zero *Nanobdellota* and zero *Patescibacteriota* tips (i.e., zero our_nano, zero our_cpr, zero r232_nano, zero r232_cpr) and pruned them, keeping 5 anchor tips per pruned clade for form-assignment context (3,888 to 1,154 tips). Both pruned trees were rooted on a Form IB plant rbcL outgroup; the analysis tree on which event enumeration is performed (`rbcl_v2.pruned.plants_rooted_trimmed.contree`) has 1,150 leaves after the four-tip plant outgroup is removed post-rooting. The pruning is applied only to the consensus tree; the underlying inference is on the full 4,262-tip alignment. Forward (archaea-to-CPR) and reverse (CPR-to-archaea) event-enumeration criteria are detailed in Supplementary Methods S1.1; pure-CPR subclade enumeration in S1.2; reproducibility pointers and limitations in S1.3 and S1.4.

### rbcL cross-kingdom protein clustering at sequence-identity sweep

To detect clusters that contain both archaeal and *Patescibacteriota* rbcL sequences from our environments, the 133 strict-K01601 hits across our 256 *Nanobdellota* and the 42 strict-K01601 hits across our 632 co-sampled high-quality *Patescibacteriota* were combined into a single 175-sequence FASTA (`rubisco_tree_v2/build_175_rbcl.py` writes `rubisco_tree_v2/combined_175_rbcl.faa`); NCBI *Nanobdellota* MAGs are excluded from this cluster sweep because they are not co-sampled with the 632 our-CPR genomes (the sweep is restricted to environmentally co-sampled archaea and bacteria). mmseqs2 v18.8cc5c easy-cluster was run repeatedly at sequence-identity thresholds of 90%, 85%, and 80% with mutual coverage 80% (`mmseqs easy-cluster combined.faa cluster_`

`<id> tmp --min-seq-id`

`<id>-c 0.80 --cov-mode 0 --cluster-mode 0`); cluster-mode 0 selects connected components in the bidirectional best-hit graph, ensuring that any cross-kingdom cluster reflects a topologically connected sequence-identity neighborhood rather than a single best-hit edge. Clusters were classified post hoc by parsing `_cluster.tsv` and tagging each member as nano/cpr by its source proteome. The single cross-kingdom cluster identified at 80% identity (one Baltic *Minisyncoccia* rbcL plus six *Nanobdellota* rbcL) was the only cluster containing both kingdoms across the three thresholds; pairwise within-cluster identity was confirmed by `mmseqs easy-search` of the seven cluster members against each other (`scripts/ cluster_identity.py`, in this paper’s `scripts/` directory).

### rbcL transposase synteny analysis

For each *Nanobdellota* and *Patescibacteriota* rbcL hit identified by KofamScan at the strict K01601 threshold, the per-genome Pyrodigal v3.6.3 GFF was scanned for predicted protein-coding genes within ±10 ORFs of the rbcL locus on either strand (`rubisco_tree_v2/ rbcl_synteny.py`). Each neighboring ORF was annotated by KofamScan strict-threshold KO assignment plus MMseqs2 v18.8cc5c easy-search against UniRef90 (sensitivity 7.5, e-value threshold 10⁻³). Transposase-family genes were identified by the union of (i) KEGG KOs whose canonical KO name contains the substring “transposase” (loaded at runtime from `/Kittens/Data/kofamscan/ko_list`; this includes K07491 (REP-associated tyrosine transposase) and other transposase KOs in the K07xxx and IS-family ranges, with K07491 the only KO from this set with strict hits in our 256 *Nanobdellota* proteomes) and (ii) UniRef90 best hits whose top description contained the substring “transposase” (case-insensitive). For the *Pacearchaeales* SA1420A_u50695000 reported in Discussion *rbcL*, a single K07491 hit was identified four ORFs downstream of rbcL on the same strand. The *Minisyncoccia* B10_u4232805 rbcL neighborhood contains no transposase-family hits within ±10 ORFs.

### rbcL directionality permutation test

To test whether the observed 9-vs-2 forward-vs-reverse event count exceeds what the source-pool size asymmetry would predict on its own, we permuted the kingdom labels (`cpr`, `nano`, `arch_other`) among the 1,132 tips carrying those labels in the 1,150-tip plants-trimmed analysis tree, leaving the 18 KEGG and outgroup-residual tips fixed; tree topology was held constant. For each of 10,000 permutations, the same forward and reverse event-enumeration logic (Methods *rbcL phylogeny*) was re-run and the per-direction event counts recorded.

Reproduction script: `rubisco_tree_v2/rbcl_permtest.py`. The null distribution recovered mean 109 forward events (5/95th percentiles 96/121; range 80–138) and mean 18 reverse events (5/95th percentiles 13/25; range 5–33), with a null mean asymmetry of 90 forward-minus-reverse events (range 54–125 across 10,000 permutations).

Interpretation in Results *Form III rbcL transfer from archaea to CPR*: the observed asymmetry is in the same direction as the null but smaller in magnitude than the source-pool sizes alone predict, so the count comparison is not a calibrated rate; the per-event phylogenetic structure (clade nesting depth, UFBoot support, donor lineage heterogeneity) is the primary support for the individual events.

### rbcS structural absence check

To verify the absence of the RuBisCO small subunit rbcS (K01602) at the structural level (independent of KofamScan sequence-level detection), four AlphaFold-DB rbcS reference structures were searched with Foldseek v10.941cd33 (van Kempen et al. 2024) against the 149,704-PDB *Nanobdellota* ESMFold structure collection. The four references span plant and cyanobacterial rbcS to capture cross-kingdom fold variation: P00873 (*Spinacia oleracea*), P04716 (*Lemna gibba*), P10795 (*Pisum sativum*), and P45686 (*Synechococcus elongatus*). Search invocation: `foldseek easy-search queries esmfold result.m8 tmp --threads 32-e 0.001 --format-output’query,target,fident,alnlen,evalue,bits,prob,alntmscore,qtmscore,t tmscore,lddt’`. The query set returned zero hits at e ≤ 10⁻³ across all four references, structurally corroborating the KofamScan-based absence of rbcS in the 256 *Nanobdellota* genomes. Reproduction script and queries: `rbcs_prk_foldseek_check/`. The same procedure was run for phosphoribulokinase PRK (K00855) using three AlphaFold-DB references (P09559 *S. oleracea*; P25697 *Arabidopsis thaliana*; P23015 *Synechocystis* sp. PCC 6803) and recovered fold-similarity hits in 43 of 256 Nano genomes (107 hits at TM-score ≥ 0.5, sequence identity 12–27%). PRK belongs to the P-loop NTPase superfamily, a fold shared with many unrelated kinases, so high TM-score with low sequence identity is consistent with non-PRK P-loop kinases rather than true PRK orthology; we therefore restrict the PRK absence claim to the sequence (KofamScan) level and do not extend it to structural absence. The Foldseek result table is at `rbcs_prk_foldseek_check/result.m8` and the per-query summary at `rbcs_prk_foldseek_check/parse_results.py` output.

The ESMFold structure collection used as the Foldseek target was predicted with ESMFold v1 (Lin et al. 2023) on the 272,391-protein “OF-era” *Nanobdellota* proteome (the Apr 15 2026 proteome that fed OrthoFinder; see `of_era_proteomes/nano_genes_all/`). 84 of 206 genomes were re-rotated after that proteome was frozen, which shifted Pyrodigal gene IDs in the current canonical proteome (`genes/`); for any analysis that consumes ESMFold output the OF-era proteome at `of_era_proteomes/nano_genes_all/` is the lookup of record. ESMFold prediction itself is sequence-identical regardless of contig rotation, so the structural search above is unaffected.

### Custom Form III rbcL HMMs for cross-kingdom homology test

Two custom HMMs were built to test whether archaeal and *Patescibacteriota* Form III rbcL sequences belong to the same protein family. Training set 1: the 133 strict K01601 hits (≥310.97 bits) across our 256 *Nanobdellota* genomes. Training set 2: the 42 strict K01601 hits across our 632 high-quality *Patescibacteriota* genomes co-sampled from the same environments. For each, training sequences were aligned with MAFFT v7.525 L-INS-i (`--localpair --maxiterate 1000`), trimmed with BMGE v2.0 under BLOSUM30, and used to build an HMM with HMMER v3.4 hmmbuild (default options). Each HMM was then searched against the opposite kingdom’s full proteome set at e<10⁻⁵ and the hit set compared to the K01601 hit set. HMM files are available in the analysis archive at `custom_hmms/rbcl/`.

### Custom Nanobdellota-trained KO HMMs for KofamScan validation

KofamScan KEGG HMMs are trained on broad protein-family alignments dominated by eukaryotic and well-characterized bacterial sequences, and we observed empirically that a substantial fraction of strict-threshold absences in our *Nanobdellota* proteomes were rescued at relaxed thresholds with UniRef90 confirmation — i.e., the gene is present but the standard KEGG HMM under-detects it. To quantify and correct this for the metabolic claims in the paper, custom *Nanobdellota*-trained HMMs were built for 154 KOs spanning core metabolism (TCA, glycolysis, gluconeogenesis, amino-acid biosynthesis, cofactor biosynthesis, pentose-phosphate pathway, RuBisCO/CBM, and information-processing). For each KO, training sequences were drawn from our 256 *Nanobdellota* proteomes by `hmmsearch` with the KEGG HMM at 0.5× the strict score threshold (the same lower bound used to define “relaxed threshold” in *Functional annotation* above) to maximise recall against divergent archaeal homologs, then filtered programmatically by retaining the highest-scoring hit per genome (multi-hit genomes were collapsed to one representative; per-tool implementation in `custom_hmms/calibrate_hmms.py`). Sequences were aligned with MAFFT v7.525 L-INS-i (`--localpair --maxiterate 1000`), trimmed with BMGE v2.0 under BLOSUM30, and used to build custom HMMs with HMMER v3.4 hmmbuild (default options). Each custom HMM was searched against the 256 *Nanobdellota* proteomes at e<10⁻⁵ and the per-KO recovery rate compared to the KofamScan strict baseline. For a per-KO verdict, recovery on *Woesearchaeales*/ *Pacearchaeales*/SCGC was extracted before and after the custom HMM was substituted, and verdicts were assigned: ROBUST (≥10 percentage point improvement, with an average paralog count <2.0; n=94), EXPANDED (≥10 percentage point improvement but average paralog count ≥2.0; n=23), COLLAPSED (no recovery improvement and the KO genuinely absent across the phylum; n=25), or PARTIAL (mixed signal at the order level; n=12). The 94 ROBUST HMMs were used to overlay marker presence on the ar71 phylogeny (Results *RuBisCO and the metabolic divide across* Nanobdellota* orders*); the per-KO verdict file is at `custom_hmms/kofam/hmm_verdicts.tsv` (column 1 of that TSV is the canonical list of all 154 KOs in the calibration set) and the HMMs themselves at `custom_hmms/kofam/`. KofamScan strict thresholds were retained for absence claims to remain conservative; custom-HMM rescues were used only to confirm presence (i.e., as a positive control on the KofamScan strict result, not as a replacement). For absences specifically of electron transport chain subunits, where under-detection is most acute, an additional set of archaea-trained HMMs was constructed (next subsection).

### Archaea-trained HMMs for validating ETC subunit absences

KEGG KofamScan HMMs are trained on sequences drawn predominantly from eukaryotes and well-characterized bacteria, and systematically under-detect divergent archaeal orthologs — we observed this repeatedly in pilot work (for example, K02603, the KEGG Cdc6/Orc1 HMM, failed to recover Cdc6 in any of 70 *Methanococcales* genomes despite Cdc6 being universal in archaea). To validate the absence of electron transport chain subunits in our *Nanobdellota* against this confounder, we tested 30 ETC-related KOs (Complex I nuoA-N, Complex II sdhABCD, Complex III petABC, Complex IV coxABC, and F-type ATP synthase atpABCDEF) and built custom archaea-trained HMMs for the 25 with at least 4 strict-threshold archaeal hits in the non-*Nanobdellota* training pool. Five KOs were excluded for insufficient training data: K00336 (nuoG), K00411 (petA), K00412 (petB), K00413 (petC), and K02115 (atpG). For comparison, K02113 (atpH) and K02114 (atpC) were retained at n=6 each. For each KO, training sequences were drawn from the 8,776 high-quality non-*Nanobdellota* archaeal proteomes in GTDB R232 via `hmmsearch` with the KEGG K-number HMM at the profile-specific strict score threshold, restricted to the top 200 hits per KO. Training sequences were aligned with MAFFT v7.525 L-INS-i, trimmed with BMGE v2.0 under BLOSUM30, and used to build custom HMMs with hmmbuild (default options). Each archaea-trained HMM was then searched against our 256 *Nanobdellota* proteomes at e<10⁻⁵. A hit in our *Nanobdellota* against the archaea-trained HMM is treated as evidence of genuine presence; absences at this more-sensitive threshold confirm the KofamScan absence. The 25 HMMs and per-KO result tables are available in the analysis archive at `custom_hmms/kofam_etc/`.

### Correlation analyses

Pearson correlations were computed in Python v3.12 (SciPy v1.14 pearsonr) for: (i) per-genome RuBisCO presence (binary) versus glycolytic gene count (number of 6 key glycolytic KOs present: K00845, K01624, K00134, K00927, K01689, K00873), n = 446; (ii) per-genome genome size versus unique KO count, n = 446. The 446-genome sample combines the 208 complete genomes from this study and the 238 NCBI genomes with KofamScan data.

### tRNA and rRNA detection

tRNAs were predicted with tRNAscan-SE v2.0.12 (Chan et al. 2021) in archaeal mode (-A, using Infernal with archaeal covariance models TRNAinf-arch.cm, default score cutoff 20 bits) on each of the 208 complete genomes. rRNA genes were detected with Infernal cmsearch v1.1.5 (Nawrocki and Eddy 2013; –tblout –noali –cpu 16, default per-sequence reporting E-value threshold of 0.01) using Rfam covariance models for SSU rRNA (RF00177), LSU rRNA (RF02541), and 5S rRNA (RF00001) on the same 208 genomes. Per-amino-acid presence was tallied from the tRNAscan-SE TSV outputs treating canonical tRNAs only (Undet, Pseudo, Sup, and iMet annotations excluded; fMet collapsed into Met).

### Frameshift detection

Assembly frameshifts were detected in the 208 complete genomes by two methods (`scripts/frameshift_detection.py`). Method 1 (split genes): adjacent same-strand ORFs within 200 bp whose predicted proteins both hit the same UniRef90 target (e-value ≤ 10⁻³ from the easy-search above). Method 2 (truncated-with-gap): genes whose predicted protein is < 70% the length of their UniRef90 target, followed by an intergenic gap (30 bp to 1.3× the missing portion in nucleotides) on the same strand; the gap was 3-frame translated, the longest open reading frame fragment (≥ 20 aa) was extracted, and cases where the combined gene + best gap fragment length was 0.7–1.5× the target length were retained (no independent database search of the gap fragment).

### Using the resource

The Zenodo deposit packages the artefacts needed to apply this study to a reader’s own *Nanobdellota* or DPANN proteomes:

- *ar71 marker set, supermatrix, and ML tree* — the 71 calibrated profile HMMs with embedded GA/NC bit-score thresholds, available both as 71 individual files (`custom_hmms/ar71/hmms/`) and as a single pre-concatenated, `hmmpress`-indexed bundle (`custom_hmms/ar71/ar71_combined.hmm`); the per-marker metadata index (`custom_hmms/ ar71/index.tsv`); the 1,239-taxon concatenated BMGE-trimmed supermatrix (`ar71_tree/supermatrix.faa`); and the inferred ML topology (`ar71_tree/ar71.treefile`). ar71 was built by an iterative HMM-refinement procedure with calibrated per-marker bit-score thresholds and a domain-coverage filter (Methods *Marker set selection and HMM construction*). Users wishing to extract the ar71 markers from new proteomes should run `hmmsearch --cut_ga` against the deposited HMMs; the GA threshold is embedded in each HMM. Adding user genomes to the supermatrix (MAFFT `--add --keeplength` per marker) and re-inferring under LG+F+R10 is the intended reuse path.
- *Nanobdellota-trained KO HMMs* — 154 verdict-classified `.hmm` profiles for KEGG orthologs (`custom_hmms/kofam/K*_nano.hmm`), of which 94 are classified ROBUST and recommended for archaeal-sensitivity KO detection where standard KEGG HMMs under-detect in DPANN proteomes. The 94 ROBUST profiles are also shipped as a single pre-concatenated, `hmmpress`-indexed bundle (`custom_hmms/ kofam/kofam_robust.hmm`) for one-shot `hmmsearch` use. The per-KO verdict file (`custom_hmms/kofam/hmm_verdicts.tsv`) documents which of the 154 KOs are ROBUST/EXPANDED/COLLAPSED/PARTIAL together with the per-HMM score threshold used in this study; the directory also retains seven calibration-leftover `.hmm` files (161 files in total) that did not enter the final 154-KO verdict set; readers should restrict reuse to the 154 listed in `hmm_verdicts.tsv`. Users running `hmmsearch` against the per-KO HMM should apply the threshold from `hmm_verdicts.tsv` to recover the same presence definition used in the paper.
- *Form III rbcL reference tree and alignment* — the 4,262-tip consensus tree (`rubisco_tree_v2/rbcl_v2.contree`), the 4,262-tip ML treefile (`rubisco_tree_v2/rbcl_v2.treefile`), the 296-position BMGE-trimmed alignment (`rubisco_tree_v2/rbcl_v2.bmge.faa`), the IQ-TREE run log (`rubisco_tree_v2/rbcl_v2.iqtree`), and a stand-alone best-model file with the LG+R10 substitution and rate parameters in IQ-TREE serialized form (`rubisco_tree_v2/ rbcl_v2.bestModel`) for direct consumption by `pplacer` / `EPA-ng`. To place new rbcL sequences on this tree, the canonical workflow is MAFFT `--add --keeplength` alignment extension against the deposited BMGE-trimmed alignment, followed by EPA-ng or pplacer using the deposited best-model file.
- *Per-step scripts* — every script that produced a paper figure or table is included, with PEP 723 inline metadata where applicable so they run under `uv run` without an explicit environment file.

The deposit is a static Zenodo archive (DOI 10.5281/zenodo.20174424) paired with the per-step scripts in this paper’s `scripts/` directory; we do not provide an interactive portal. The resource is intended to be applied by readers in their own compute environment using the canonical bioinformatics tools cited in Methods.

## Data Availability

Raw Oxford Nanopore PromethION reads from the six metagenome samples used in this study are deposited under two NCBI BioProjects. `PRJNA1461169` (Fennoscandian groundwater) covers the two Äspö samples KR0015B (BioSample `SAMN57747384`) and SA1420A (`SAMN57747385`). `PRJNA1461195` (Baltic Sea water column) covers the four Baltic samples used here: B5 (`SAMN57849835`), B7 (`SAMN57849837`), B10 (`SAMN57849840`), and B14 (`SAMN57849843`). SRA-run accessions for the per-flowcell fastq files are linked under each metagenome BioSample. The 256 *Nanobdellota* genomes (208 complete + 48 HQ non-circular) have been deposited as third-party annotated GenBank entries under the same two BioProjects, with one BioSample per genome; per-genome BioSample and GenBank accessions are linked from each BioProject page. The full analysis archive — alignments, gene-tree sets, supermatrices, custom HMM archive, all per-step scripts — is deposited at Zenodo under DOI 10.5281/zenodo.20174424. A standalone Supplementary Information document (`supplementary_information.md`) accompanies this paper and contains Supplementary Table S1 (CPR-to-archaea reverse-direction candidate events), Supplementary Table S2 (101-OG calibration outcomes for the ar71 marker set), Supplementary Note S1 (pure-CPR subclade backdrop in the rbcL gene tree), and Supplementary Methods (event enumeration criteria, reproducibility pointers, limitations).

## Acknowledgments

We thank Stefan Bertilsson for providing the Baltic Sea water-column samples (B5, B7, B10, B14); Mark Dopson for providing the Fennoscandian groundwater samples (KR0015B, SA1420A); and the Swedish Nuclear Fuel and Waste Management Company (SKB) for borehole access. This work was partially supported by the Laboratory Directed Research and Development Program of Lawrence Berkeley National Laboratory under the U.S. Department of Energy (DOE) (Contract No. DE-AC02-05CH11231).

## Author contributions

Both authors contributed equally to all aspects of this study, including study design, data analysis, and manuscript preparation.

## Competing interests

The authors declare no competing interests.

---

|  | Complete (this study) | NCBI MAGs | Combined |
| --- | --- | --- | --- |
| *Pacearchaeales* | 100% (54/54) | 69% (20/29) | 89% (74/83) |
| *Maxwellarchaeales* | 87% (27/31) | 77% (34/44) | 81% (61/75) |
| *Woesearchaeales* | 11% (14/123) | 10% (8/82) | 11% (22/205) |

Lower NCBI rates in *Pacearchaeales* and *Maxwellarchaeales* are consistent with MAG incompleteness; identical rates in *Woesearchaeales* suggest the low Woese prevalence is genuine. No order has a rbcS hit at either sequence (KofamScan) or structural (Foldseek vs AlphaFold-DB) level, and no order has a KofamScan ortholog call for PRK, confirming Form III identity and ruling out the Calvin cycle.

## Notes

### Competing Interest Statement

The authors have declared no competing interest.

