## Supplemental Tables for "A complete-genome view of phylum Nanobdellota and recurrent Form III RuBisCO transfer between archaea and Patescibacteriota"

Nielsen TN, Lui LM

---

#### Supplementary Table S1. Reverse-direction (CPR→archaea) candidate events in the pruned v2 rbcL gene tree

Mirror of the Table 3 analysis with CPR and archaea roles inverted. The same topological criteria were applied (clade size ≤21 tips, ≥3:1 majority) but with CPR as the majority and Nanobdellota rbcL as the nested minority tip(s). Two such candidate events were identified, involving three Nanobdellota rbcL tips in total; both candidate events involve R232 HQ Nanobdellota rbcL rather than rbcL from our 206 complete genomes.

| Event | Clade size | CPR tips | Nano tips | UFBoot | Nanobdellota rbcL tip(s) |
|---|---|---|---|---|---|
| S1.1 | 12 | 3 | 1 | 57 | r232_nano GCA_903825585.1 |
| S1.2 | 17 | 6 | 2 | 41 | r232_nano GCA_023266675.1; one additional R232 Nano sister tip |

UFBoot values were extracted from the pruned plants-rooted v2 contree at each event's defining MRCA by `rubisco_tree_v2/event_ufboot.py`. Both reverse-direction candidate events fall below the UFBoot ≥ 80 confidence threshold; under that threshold, the reverse direction contributes zero qualifying events while the forward direction contributes five (Events 1, 2, 3, 5, 9 at UFBoot 96, 97, 99, 97, 81 respectively), giving a 5:0 directional ratio in the higher-confidence subset.

For comparison, the forward (archaea→CPR) direction yielded 9 events involving 12 CPR tips at the same criteria (Table 3), representing a 4.5:1 asymmetry by event count and 4:1 asymmetry by tip count under the unfiltered ≤21-tip / ≥3:1 majority criterion. Neither of the two reverse-direction candidate events involves our 206 complete Nanobdellota genomes.

### Supplementary Table S2. Calibration outcomes for the 101 single-copy orthogroups and selection of the 71-marker ar71 set

For each of the 101 OrthoFinder single-copy orthogroups inferred across the 206 complete genomes, a profile HMM was built and iteratively refined against a 1,288-genome calibration pool (the 256 Nanobdellota proteomes from this study plus 1,031 GTDB R232 high-quality Nanobdellota MAGs and the 238 HQ NCBI Nanobdellota MAGs), with per-marker GA/NC bit-score thresholds calibrated against 206-genome OrthoFinder OG-membership ground truth (Methods §Marker set selection and HMM construction). Each orthogroup was evaluated on three quality criteria: single-copy recovery on the calibration pool ≥1,100/1,288, multi-copy rate ≤5%, and GA value ≥30 bits. **71 of 101 orthogroups satisfied all three criteria** and constitute the ar71 marker set; the remaining 30 failed for paralog distribution across Nanobdellota subsets, under-detection in distantly-related orders, or score non-separability. Per-marker overlap with the standard GTDB archaeal marker sets ar53 and ar122 is recorded in `custom_hmms/ar71/index.tsv`. The full per-OG calibration table — with sc/mc counts, GA/NC values, and pass/fail verdict per orthogroup — is at `custom_hmms/ar71/index.tsv` together with the calibrated HMMs (`custom_hmms/ar71/hmms/`).

### Supplementary Note S1. Pure-CPR subclade background in the pruned v2 rbcL gene tree

To contextualize the 9 archaea→CPR + 2 reverse-direction events, we counted subclades in the pruned v2 tree (1,154 tips post-Strategy-A/B reduction; 1,150 tips after the 4-tip plant Form IB outgroup is removed post-rooting on the analysis tree `rbcl_v2.pruned.plants_rooted_trimmed.contree`; see Methods) whose composition is ≥90% Patescibacteriota rbcL and contain zero Nanobdellota rbcL tips, at subclade sizes 10 to 200 tips. 66 such pure-CPR subclades were identified. This indicates that CPR carries extensive vertically-inherited Form III rbcL beyond what is visible at our sampling scale: the 9 archaea→CPR transfer events affect 12 CPR rbcL tips out of 320 in the v2 analysis (~4%), while the remaining ~96% of CPR rbcL occupies the 66 pure-CPR subclades and smaller pure-CPR cherries. The archaea→CPR transfers we identify in Table 3 are interlopers into an otherwise taxonomically coherent CPR Form III subtree.

### Supplementary Methods

#### S1.1 Criteria for archaea→CPR and CPR→archaea event enumeration

Both directional analyses used the same deep-nesting criterion: a candidate event is a CPR (or Nanobdellota) rbcL tip whose smallest ancestral clade with size ≥4 and ≤21 tips contains a majority of the opposite kingdom at ≥3:1 ratio. The lower bound of 4 is the minimum size that admits the ≥3:1 ratio with at least one focal-kingdom tip (3 of one kingdom + 1 of the other). Clades were counted under the pruned v2 tree (1,154 tips post-Strategy-B; 1,150 leaves after outgroup trim) rooted on the 4-tip plant Form IB cherry (`kegg|cmax_111494870`, `kegg|csat_109132455`, `kegg|rcu_107261263`, `kegg|bhw_143583404`) with outgroup tips dropped post-rooting. Branch lengths were not used in the enumeration — only topology and tip identity.

#### S1.2 Pure-CPR subclade enumeration

For every internal node in the pruned and rooted v2 tree, we recorded composition as (n_nano, n_cpr, n_other). A node was classified "pure-CPR" if size was between 10 and 200 tips, n_cpr was ≥90% of size, and n_nano was zero. Overlapping ancestor–descendant subclades were retained as separate counts to give a non-deduplicated structural measure of CPR subclade density; this count is therefore an upper bound on distinct Form III CPR lineages.

#### S1.3 Reproducibility

All analyses are available as standalone scripts in `rubisco_tree_v2/prune_rbcl_v2.py` (tree reduction and rooting pipeline) with inline analysis code for event enumeration. The pruned and rooted tree used for the tabulation is `rubisco_tree_v2/rbcl_v2.pruned.plants_rooted_trimmed.contree`. UFBoot support values are preserved on internal nodes of this tree for downstream robustness assessment.

#### S1.4 Known limitations

1. UFBoot correlation of the underlying v2 tree was 0.922 at run completion, below the 0.99 convergence threshold. Event boundaries can shift between consecutive bootstrap replicates; our numerical results (9 archaea→CPR events, 2 CPR→archaea events, 66 pure-CPR subclades) are therefore point estimates rather than confidence-bounded quantities.
2. The tree was pruned from 4,262 to 1,154 tips via Strategy A (dereplicate by genome or KEGG species; 4,262 → 3,888 tips) and Strategy B (drop target-free clades ≥30 tips, keeping 5 anchors; 3,888 → 1,154 tips); after rooting on the 4-tip plant Form IB outgroup the analysis tree retains 1,150 leaves. Some CPR or Nanobdellota tips may have been lost at Strategy A when a shorter paralog won the dereplication; no evidence of this affecting event boundaries was found in our inspection, but it cannot be ruled out.
3. The plant Form IB outgroup is a 4-tip dicot cherry (`cmax`, `csat`, `rcu`, `bhw`) rather than a curated Form I reference set; bacterial Form I/II/IV references were deliberately excluded from v2 per the clean-sources policy. Sensitivity of event counts to alternative rooting was not tested quantitatively here; however, the topological nesting patterns that define each event are largely rooting-independent once a consistent outgroup position is selected.
